# Comparative characterisation of *Plasmodium falciparum* Hsp70-1 relative to *E. coli* DnaK reveals functional specificity of the parasite chaperone

**DOI:** 10.1101/2020.03.09.984104

**Authors:** Charity Mekgwa Lebepe, Pearl Rutendo Matambanadzo, Xolani Henry Makhoba, Ikechukwu Achilonu, Tawanda Zininga, Addmore Shonhai

## Abstract

Hsp70 is one of the most prominent molecular chaperones. Although Hsp70s from various organisms are generally conserved, they exhibit specialised cellular functions. It remains to be fully understood how these highly conserved molecules exhibit specialised functional features. *Plasmodium falciparum* Hsp70-1 (PfHsp70-1) is a cytosol localised molecular chaperone that is implicated in the cyto-protection and pathogenicity of the malaria parasite. In the current study, we investigated the comparative structure-function features of PfHsp70-1 relative to its homologue, *E. coli* Hsp70 (DnaK) and a chimeric protein, KPf, that was constituted by the ATPase domain of DnaK and the substrate binding domain (SBD) of PfHsp70-1. Recombinant forms of all the three Hsp70s exhibited similar secondary and tertiary structural fold. We further established that compared to DnaK, both KPf and PfHsp70-1 were more stable to heat stress and exhibited higher basal ATPase activity. A recombinant *P. falciparum* Hsp40 (PfHsp40) stimulated the ATPase activities of all the three Hsp70s. In addition, both PfHsp70-1 and KPf exhibited preference for asparagine rich peptides as opposed to DnaK. Furthermore, all the three proteins exhibited self-association capabilities in vitro. Recombinant *P. falciparum* adenosylmethionine decarboxylase (PfAdoMetDC) co-expressed in *E. coli* with either KPf or PfHsp70-1 was produced as a fully folded product. On the other hand, co-expression of PfAdoMetDC with heterologous DnaK in *E. coli* did not promote folding of the former. These findings demonstrated that the SBD of PfHsp70-1 regulates several functional features of the protein and that this molecular chaperone is tailored to facilitate folding of plasmodial proteins.

---

Heat shock protein 70 (Hsp70) molecular chaperones are involved in protein folding, unfolding, assembly and disassembly of protein units and they also facilitate protein degradation. *Plasmodium falciparum* Hsp70-1 (PfHsp70-1) is a cytosol-localised molecular chaperone that is essential for survival of the malaria parasite (1-2). PfHsp70-1 has been proposed as a prospective antimalarial drug target (2-4). Furthermore, PfHsp70-1 is implicated in antimalarial drug resistance, making its inhibition using antimalarial drug combinations promising (5). Although some compounds that inhibit PfHsp70-1 have been reported to be selective, exhibiting minimum effects on human Hsp70 (4), The unique structure-function features of this protein remain to be fully explored. This is important towards understanding its role in the survival of the parasite and validation of its utility as a possible antimalarial drug target. Notably, PfHsp70-1 and its homologues of parasitic origin are endowed with an extended GGMP repeat motif that is located in the C-terminal substrate binding domain (SBD) (6). The location of the GGMP motif in the SBD of PfHsp70-1 suggests possible roles of this motif in regulating substrate selection and interaction of the chaperone with its co-chaperones.

Structurally, Hsp70 consists of two functional domains: an N-terminal nucleotide-binding domain (NBD) and a C-terminal SBD (7). The NBD of Hsp70 binds to ATP hydrolysing it to ADP (8). On the other hand, the SBD binds to the peptide substrate. The SBD of Hsp70 is sub-divided into α- and β-subdomains. The Hsp70 SBDβ makes direct contact with the peptide substrates while the SBDα serves as a lid enclosing the bound substrate (9). Hsp70 exhibits high affinity for substrate in its ADP-bound state and releases it upon binding to ATP (7). Therefore, nucleotide binding regulates substrate binding and release by the Hsp70 chaperone. Hsp70 possesses weak basal ATPase activity and hence relies on a co-chaperone, Hsp40 that stimulates its ATPase activity (10). In addition, Hsp40 binds to misfolded proteins first before transferring them to Hsp70 for refolding (11). Thus, delivery of substrate to Hsp70 is concomitantly linked to ATP hydrolysis (12). Nucleotide exchange on Hsp70 is facilitated by nucleotide exchange factors (NEFs) represented by GrpE in *E. coli* (10).

*E. coli* Hsp70 (DnaK) structurally constitutes a canonical Hsp70, characterised by a conserved NBD connected to the SBD via a highly conserved linker motif (13). Although DnaK is not essential for *E. coli* growth at intermediate temperatures, cells lacking DnaK die at high temperatures (14). PfHsp70-1 is a homologue of DnaK that localises to the malaria parasite cytosol (15). PfHsp70-1 and a chimeric protein, KPf (made up of the ATPase domain of DnaK and the SBD of PfHsp70-1) were previously shown to protect *E. coli* dnaK756 cells (harbouring a functionally compromised DnaK), against heat stress (16). In addition, PfHsp70-1 has been shown to provide cyto-protection to yeast cells endowed with a defective Hsp70 (17). Altogether, this suggests that although Hsp70s are functionally specialised they also exhibit functional overlaps across species.

PfHsp70-1 is stress-inducible, and its inhibition leads to parasite death suggesting its essential role in parasite cyto-protection (18, 19). It was previously reported that PfHsp70-1 possesses higher basal ATPase activity compared to its human and *E. coli* Hsp70 homologues (3). The unique structure-function features of PfHsp70-1 in comparison to its human Hsp70, have spurred interest to target it as part of anti-malarial drug design efforts (3, 4, 6). *P. falciparum* Hsp40 (PF3D7-1437900) is a co-chaperone of PfHsp70-1 that co-localises with it to the parasite cytosol (20). PfHsp40 is regarded as a member of the so-called type I Hsp40s (on account its structural resemblance to *E. coli* Hsp40 (DnaJ; 21). Both DnaJ and PfHsp40 possess a highly conserved J domain that facilitates cross-talk with Hsp70 (21). It has been shown that Hsp40 chimeric proteins with J domains of variable eukaryotic origin were able to cooperate with DnaK to confer cyto-protection to *E. coli* cells that were void of endogenous DnaJ (22). This suggests that the highly conserved J domain of Hsp40 is capable of modulating the function of Hsp70s of varied species origin. Indeed, functional interaction between Hsp40 and Hsp70 of varied species origin has been demonstrated experimentally (23), further highlighting that the conserved J domain mediates promiscuous Hsp70-Hsp40 partnerships.

Although functional specificity of Hsp70s across species is generally regarded to be on account of their cooperation with several Hsp40 partners with whom they interact (24), we still do not understand how such conserved molecules are adapted to their function. It is further believed that of the two domains of Hsp70, it is the less conserved SBD that provides it with functional specificity (8). For example, Hsp70s of eukaryotic origin are thought to possess arch residues (located in their substrate binding cavity) that are inverted in orientation in comparison to those of prokaryotes (8, 25). The arch residues are thought to regulate functional specificity of Hsp70 as they constitute part of the residues that make direct contact with the substrate (8).

It has been proposed that nearly 10% of *P. falciparum* proteome is characterized by prion-like repeats and that at least 30% of the proteome is characterized by glutamate/asparagine rich segments (26, 27). For this reason, it is thought that *P. falciparum* Hsp70s are adapted to fold its aberrant-prone proteome (28, 29, 30). To this end, we previously demonstrated that a *P. falciparum* chaperone, PfHsp70-x, that is exported to the parasite-infected red blood cell (31) exhibits preference for asparagine rich peptides, further suggesting that *P. falciparum* Hsp70s are primed to bind misfolded proteins of the parasite (30).

Both PfHsp70-1 and its chimeric product, KPf have been shown to confer cyto-protection to *E. coli* cells harbouring functionally compromised DnaK (15). This suggests that PfHsp70-1 and KPf exhibit functional overlap with DnaK. On the other hand, PfHsp70-1 and KPf have both been employed to improve the quality and yield of recombinant proteins of plasmodial origin expressed in *E. coli* (29, 32). This suggests that although PfHsp70-1 and KPf exhibit functional overlap with DnaK, they are particularly tailored to facilitate folding of proteins of plasmodial origin. For these reasons, PfHsp70-1, DnaK and their chimeric protein, KPf, present a convenient model for studying the functional specificity of PfHsp70-1.

*P. falciparum* adenosylmethionine decarboxylase (PfAdoMetDC) is an essential protein involved in the biosynthesis of polyamines, making it a potential anti-malarial drug target (33). Previously, we demonstrated that recombinant PfAdoMetDC co-expressed in *E. coli* with either KPf or PfHsp70-1 exhibited higher enzymatic activity than that co-expressed with supplementary *E. coli* DnaK (29).

GroEL and its cofactor, GroES constitute a chaperonin of *E. coli* that is constituted of a cylindrical complex of two heptameric rings (34). Thus GroEL/ES system constitutes a cage in which some misfolded proteins are sequestered into to facilitate their folding. It has been proposed that the GroEL/ES cage accommodates substrates of up to 60 kDa in size (35). GroEL/ES and DnaK cooperate to facilitate folding of some proteins in *E. coli* (35). For this reason, we investigated the effect of the three Hsp70s on the folding status of recombinant PfAdoMetDC expressed in *E. coli* BL21 Star^™^ (DE3) cells. We further expressed each of the Hsp70 along with GroEL towards exploring their combined influence on PfAdoMetDC folding.

Our findings established that all the three Hsp70s exhibited comparable secondary and tertiary structures and they also shared some functional features. However, both PfHsp70-1 and KPf preferentially bound to peptide substrates that had enriched asparagine residues while the presence of asparagine did not enhance the affinity of DnaK for these peptides. In addition, both PfHsp70-1 and KPf were marginally more stable to heat stress than DnaK.

Our findings highlight the importance of the SBD of Hsp70 in stabilizing the conformation of this chaperone and its role in defining the functional specificity of the molecular chaperone. In addition, PfAdoMetDC co-expressed in *E. coli* with PfHsp70-1 and KPf exhibited similar biophysical features and was better folded than the protein co-produced with supplementary *E. coli* DnaK. This further demonstrates that the SBD of PfHsp70-1 is tailored to fold proteins of plasmodial origin.

## RESULTS

### Secondary and tertiary structural analysis of DnaK, KPf and PfHsp70-1

The comparative secondary structure analysis of purified recombinant forms of DnaK, PfHsp70-1 and the chimeric KPf (Fig. S1) was assessed using CD spectroscopy. The far-UV CD spectra exhibited negative troughs at 208-210 nm and 220-225 nm, representing the predominantly α-helical composition for all the three proteins (Fig. 1a). A positive peak was also observed at 190 nm, representing the β-pleated sheets of the protein as previously reported for PfHsp70-1 (36, 37). The secondary structure of each protein upon exposure to variable temperature conditions (19 °C to 95 °C) was assessed (Fig. 1b). PfHsp70-1 and KPf maintained at least 50 % fold at temperatures up to 60 °C. However, DnaK rapidly unfolded at temperatures above 45 °C and completely lost its fold at approximately 68 °C. On the other hand, KPf and PfHsp70-1 became completely unfolded at around 90 °C. Our findings suggest that KPf and PfHsp70-1 were nearly equally resilient to heat stress at temperatures below 60 °C and this is in agreement with a previous study (37), which reported that PfHsp70-1 is stable at temperatures above 50 °C. Notably, both KPf and PfHsp70-1 displayed a unique unfolding pattern, characterised by two steps of which the second phases occurred at temperatures 55 °C – 90 °C for KPf; and 75 °C - 90 °C for PfHsp70-1, respectively.

**Figure 1.**
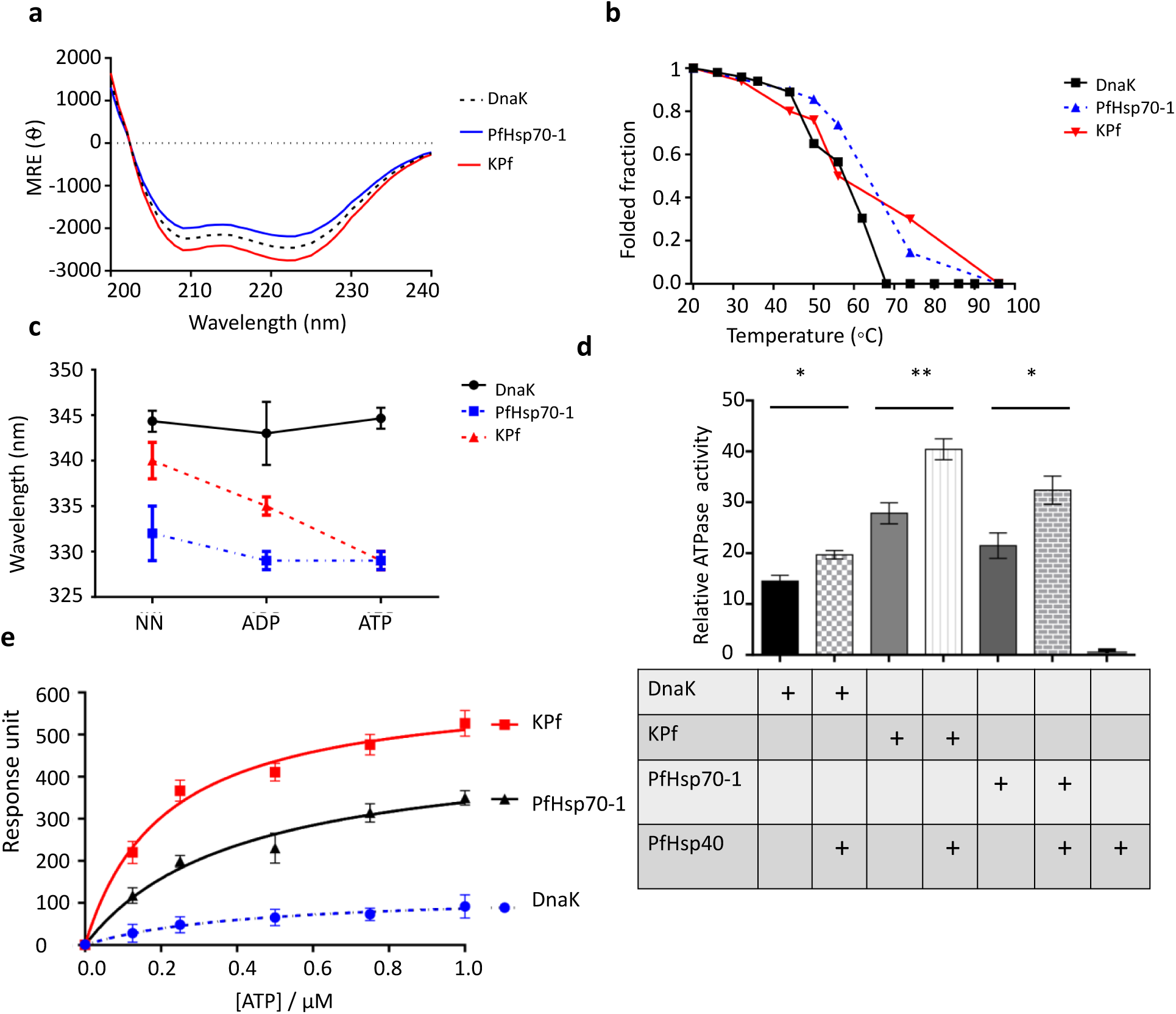
Comparative analysis of heat stability, nucleotide binding and hydrolysis kinetics of the Hsp70s. **(a)** *Hsp70 far-UV spectra*. The far-UV CD spectra of recombinant DnaK, KPf and PfHsp70-1 proteins were presented as molar residue ellipticity. All the three Hsp70s displayed a predominantly α-helical structure due to the negative troughs observed at 209 and 222 nm. **(b)** *Comparative secondary structural fold of DnaK, PfHsp70-1 and KPf exposed to heat stress*. The folded fraction of each protein was determined by comparing the spectral readings obtained at any given temperature with readings taken at 19 °C. **(c)** *ATP dramatically modulates the conformation of KPf*. Fluorescence emission spectra were monitored at 320-450 nm after an initial excitation at 295 nm. The tryptophan fluorescence emission spectra maxima were recorded for DnaK, PfHsp70-1 and KPf proteins in the absence and presence of 5 mM ATP/ADP. The fluorescence maxima for each protein was blue shifted in the presence of either ATP or ADP. (**c**) *ATPase activities of PfHsp70, KPf and DnaK*. Inorganic phosphate released by ATP in the presence of each Hsp70 was monitored by direct calorimetry at 595 nm wavelength. The observed basal- and PfHsp40-stimulated ATPase activities of respective protein were illustrated as bar graphs. The independent activity of PfHsp40 was used as a non-ATPase control. The statistical significance of the relative ATPase activities of the proteins are shown; *p* <*0.05*\* and *p*<*0.01*\*\* as determined by ANOVA. (**e**) *Equilibrium ATP binding kinetics of all three Hsp70s*. The reported ATP binding constant for the respective Hsp70 was determined at equilibrium. Standard deviations shown represent three independent assays made using separate protein purification batches. The data was expressed as mean (±) standard deviation.

The tertiary structural conformations of DnaK, KPf and PfHsp70-1 were determined using tryptophan fluorescence either in the absence or presence 5 mM ATP/ADP. PfHsp70-1 has three tryptophan residues (W32, W101 and W593); while KPf possesses two tryptophan residues (W102 and W578). On the other hand, DnaK possesses a single, W291 residue. Due to the varying tryptophan residue composition of the proteins, we focussed on comparing the emission peaks rather than fluorescence quantum yields of the proteins (Fig. 1c). DnaK gave an emission peak at 345 nm and a marginal blue shift was observed in the presence of nucleotides (5 mM ATP/ADP) (Fig. 1c). Similarly, PfHsp70-1 displayed an emission peak at 333 nm and exhibited a marginal blue shift in the presence of ATP/ADP (Fig. 1c). On the other hand, KPf had an emission peak at 340 nm, and generated a significant blue shift in the presence of nucleotides (emission peaks of 335 nm in the presence of ADP; and 330 nm in the presence of ATP; Fig. 1c). These data suggest that KPf exhibited a tertiary conformation that slightly varied to its parental forms. Furthermore, ATP had the most marked effect on the tertiary conformation of KPf than it had on DnaK and PfHsp70-1.

### PfHsp40 stimulates the ATPase activities of all three Hsp70s

The capability of DnaK, PfHsp70-1 and KPf to hydrolyse ATP independently was determined using a calorimetric assay (Fig. 1d). The basal ATPase activity of DnaK was the lowest, while that of PfHsp70-1 was higher (*p = 0.005*) than that of DnaK. On the other hand, KPf registered the highest basal ATPase activity (Fig. 1d; Table S1). Next, we sought to explore the stimulatory effect of PfHsp40 on the ATPase activities of all the three Hsp70s. First we validated that the PfHsp40 protein preparation had no detectable independent basal ATPase activity as expected. As has been previously reported, PfHsp40 stimulated the ATPase activity of PfHsp70-1 (20). The highest ATPase activity was registered by KPf in the presence of PfHsp40 (Fig. 1d). On the other hand, the ATPase activity of DnaK was only marginally enhanced in the presence of PfHsp40. The chimeric protein, KPf, exhibited the highest ATPase activity (both basal and PfHsp40 stimulated). While Hsp40 primarily binds to the NBD domain to stimulate ATP hydrolysis by Hsp70, it is also known to make contact with the C-terminus of Hsp70 (38). It is thus possible that the contact that PfHsp40 makes with the C-terminus of the SBD of PfHsp70-1 (also present in KPf) is unique compared to that of DnaK. This may account for the comparatively higher ATPase activities that both KPf and PfHsp70-1 register in the presence of PfHsp40. Whereas both PfHsp70-1 and KPf possess C-terminal EEVD residues known to make direct contact with Hsp40 (39), the equivalent C-terminal segment in DnaK is represented by residues, EEVKDKK.

### KPf exhibits marginally higher affinity for ATP than either DnaK or PfHsp70-1

The relative nucleotide binding affinities of KPf, PfHsp70-1 and DnaK, were determined (Fig. 1e). KPf exhibited a *K*_*D*_ value which was approximately one order of magnitude higher than that of PfHsp70-1 and at least 200-fold higher than that of DnaK (Table S2). Since KPf and DnaK share the same NBD the expectation would be that these two proteins exhibit comparable affinity for ATP. However, it is known that ATP binding at the NBD allosterically modulates Hsp70 to assume a closed conformation in which the C-terminal segment, in particular the lid section, comes into contact with the NBD and this contributes towards ATP hydrolysis (40, 41). For this reason, the NBD cooperates with the SBD to influence both ATP binding and hydrolysis. This may explain why KPf exhibits much higher affinity for ATP than DnaK, despite the two proteins sharing the same NBD. Such a scenario highlights the role of the SBD:NBD interface in regulating ATP binding. The high affinity that KPf has for ATP could partly account for the marked blue shift in the tryptophan fluorescence signal that KPf displayed in the presence of ATP (Fig. 1e).

### The chimeric Hsp70, KPf is capable of self-association

The self-association of Hsp70 is thought to be mediated by both the interdomain linker and the SBD (42, 43). For this reason, using SPR analysis, we investigated the capability of KPf to self-associate relative to its parental forms, DnaK and PfHsp70-1. The immobilized protein preparation served as ligand while the protein in solution was used as the analyte. Our findings established that all the three proteins were capable of self-association (Table 1). In addition, the oligomerisation of the Hsp70s occurred in the absence of nucleotide as well as in the presence of 5 mM ADP/ATP (Table 1). However, the oligomerisation of both PfHsp70-1 and KPf occurred with higher affinity in the presence of ATP than in the ADP or absence of nucleotides. On the other hand, in the presence of ATP, the affinity for DnaK self-association was one magnitude lower than that of either KPf or PfHsp70-1 (Table 1). Overall, our findings suggest that KPf, DnaK and Hsp70-1 are all capable of forming higher order oligomers and that ATP enhances this process. Independent studies previously reported that DnaK forms higher order oligomers (44, 45). It is interesting to note that the fusion protein, KPf, retained this important functional regulatory feature of Hsp70 in vitro.

**Table 1.** Kinetics for self-association of DnaK, KPf and PfHsp70-1.

| Ligand | Analyte | $K_a$ (Ms<sup>-1</sup>) | $K_d$ (l<sup>-1</sup>) | $K_D$ (M) | $\chi^2$ |
| --- | --- | --- | --- | --- | --- |
| DnaK | DnaK+ATP | 1.38 (± 0.08) e<sup>2</sup> | 3.83 (± 0.33) e<sup>-4</sup> | 4.13 (± 0.30) e<sup>-6*</sup> | 5.15 |
|  | DnaK | 1.27 (± 0.07) e<sup>3</sup> | 6.51 (± 0.01) e<sup>-2</sup> | 4.67 (± 0.27) e<sup>-5</sup> | 4.21 |
|  | DnaK+ADP | 1.22 (± 0.13) e<sup>2</sup> | 6.49 (± 0.09) e<sup>-3</sup> | 4.60 (± 0.09) e<sup>-5</sup> | 5.41 |
| Kpf | Kpf+ATP | 1.54 (± 0.04) e<sup>2</sup> | 5.44 (± 0.04) e<sup>-3</sup> | 3.23 (± 0.03) e<sup>-7*</sup> | 2.30 |
|  | Kpf | 1.32 (± 0.02) e<sup>2</sup> | 4.17 (± 0.17) e<sup>-5</sup> | 5.38 (± 0.08) e<sup>-6</sup> | 3.17 |
|  | Kpf+ADP | 1.42 (± 0.02) e<sup>2</sup> | 5.23 (± 0.03) e<sup>-3</sup> | 4.65 (± 1.5) e<sup>-5</sup> | 2.37 |
| PfHsp70-1 | PfHsp70-1+ATP | 2.14 (± 0.04) e<sup>4</sup> | 1.13 (± 1.2) e<sup>-2</sup> | 5.28 (± 0.08) e<sup>-7*</sup> | 4.42 |
|  | PfHsp70-1 | 8.51 (± 1.20) e<sup>3</sup> | 2.04(± 0.11) e<sup>-3</sup> | 2.39 (± 0.09) e<sup>-6</sup> | 1.20 |
|  | PfHsp70-1+ADP | 1.00 (± 0.17) e<sup>2</sup> | 1.71 (± 0.07) e<sup>-4</sup> | 1.71 (± 0.10) e<sup>-6</sup> | 4.45 |

The table shows the binding kinetics parameters of self-association of the three Hsp70s. The ligand fraction of each protein was the immobilized protein on the GLC chip surface, and the analyte fraction was injected at a flow rate of 50 µl/min. Self-association of the three proteins was investigated either in the absence or presence of ATP/ADP. Significant variation in $K_D$ values was based on one way ANOVA test ( $p < 0.05$ \*).

### All the three Hsp70s directly interacted with PfHsp40

PfHsp40 is a type I (structurally resembles E. coli DnaJ) co-chaperone of PfHsp70-1 (20). In light of its high sequence homology to DnaJ and its possession of a highly conserved J domain, PfHsp40 represents a typical Hsp40 whose propensity to interact with Hsp70s from various species such as plasmodial and human Hsp70 has been demonstrated in vitro (20). In addition, in the current study we demonstrated that PfHsp40 stimulated the ATPase activities of DnaK, KPf and PfHsp70-1, further confirming its functional versatility (Fig 1d). We further sought to establish the interaction kinetics of PfHsp40 with DnaK, KPf, and PfHsp70-1, respectively. Using SPR analysis, we established that PfHsp40 directly binds to DnaK, PfHsp70-1 and KPf, respectively (Table 2). Furthermore, the interaction was enhanced in the presence of ATP, in line with a previous independent study that demonstrated that Hsp70 association with Hsp40 is favoured in the ATP-bound state of Hsp70 (46).

**Table 2.** Kinetics for the interaction of DnaK/KPf/PfHsp70-1 with PfHsp40.

|  |  |  |  |  |  |
| --- | --- | --- | --- | --- | --- |
| PfHsp70-1 | PfHsp40 +ATP | $1.81 (\pm 0.01) e^2$ | $1.32 (\pm 0.02) e^{-4}$ | $2.08 (\pm 0.80) e^{-7**}$ | 2.12 |
| | PfHsp40 +ADP | $1.53 (\pm 0.03) e^3$ | $8.66 (\pm 0.06) e^{-6}$ | $9.98 (\pm 0.18) e^{-6}$ | 3.08 |
| | PfHsp40 | $1.33 (\pm 0.03) e^3$ | $9.81(\pm 0.10) e^{-6}$ | $1.88 (\pm 0.08) e^{-5}$ | 7.80 |
| KPf | PfHsp40 +ATP | $1.72 (\pm 0.02) e^2$ | $7.44 (\pm 0.04) e^{-3}$ | $7.23 (\pm 0.30) e^{-7**}$ | 1.75 |
| | PfHsp40+ADP | $1.36 (\pm 0.26) e^2$ | $5.43 (\pm 0.03) e^{-4}$ | $2.62 (\pm 0.02) e^{-6}$ | 2.15 |
| | PfHsp40 | $1.24 (\pm 0.04) e^4$ | $6.54 (\pm 0.04) e^{-5}$ | $2.53 (\pm 0.03) e^{-5}$ | 2.13 |
| DnaK | PfHsp40 +ATP | $1.22 (\pm 0.02) e^2$ | $7.44 (\pm 0.04) e^{-4}$ | $7.23 (\pm 0.03) e^{-6*}$ | 1.66 |
| | PfHsp40 +ADP | $1.16 (\pm 0.06) e^2$ | $5.43 (\pm 0.32) e^{-3}$ | $2.62 (\pm 0.02) e^{-5}$ | 1.57 |
| | PfHsp40 | $1.12 (\pm 0.02) e^2$ | $7.44 (\pm 0.04) e^{-3}$ | $7.23 (\pm 0.03) e^{-5}$ | 2.07 |

The data was analysed through global fitting of the sensorgrams obtained. The standard deviation about the means were obtained after 3 experimental repeat runs. Statistical significance of the data was established by one way ANOVA test ( $p < 0.05$ \*) and ( $p < 0.01$ \*\*).

### PfHsp70-1 and KPf preferentially bind to asparagine-enriched peptide substrates

Approximately 10% of the malaria parasite proteome is thought to be characterized by prion-like repeats and more than 30% of the proteome is characterized by glutamate/asparagine repeat segments (26, 27). Since PfHsp70-1 and KPf both possess an SBD of plasmodial origin and have previously been shown to support expression of recombinant plasmodial proteins in *E. coli* (29, 32), we explored the comparative model Hsp70 peptide binding activities of KPf and PfHsp70-1 relative to DnaK. Thus, the binding kinetics of DnaK, KPf and PfHsp70-1 for a battery of synthetic peptide substrates was determined using SPR analysis (Table S3). The relative affinities of the respective Hsp70s for the canonical Hsp70 peptide substrates (NRLLTG, ALLLMYRR, GFRVVLMYRF) (7, 30, 47, 48) were determined by SPR analysis. The assay was repeated using peptides that were modified by substituting the middle residues with asparagine residues (NR**NN**TG,A**NNN**MYRR, GFR**NNN**MYRF; 30). DnaK bound to the standard peptides (NRLLTG, ALLLMYRR and GFRVVLMYRF) with higher affinity signals than it exhibited for the respective N-enriched peptides (Fig. 2; Table S3). On the other hand, PfHsp70-1 and KPf bound preferentially to the asparagine (N)-enriched peptides; A**NNN**MYRR and GFR**NNN**MYRF and exhibited less affinity for the original18m peptides (Fig. 2; Table S3). We previously observed that another plasmodial Hsp70, PfHsp70-x preferentially binds to asparagine rich peptides (30). Altogether these observations suggest that SBDs of Hsp70s of *P. falciparum* origin are tailored to bind to asparagine rich peptides.

**Figure 2.**
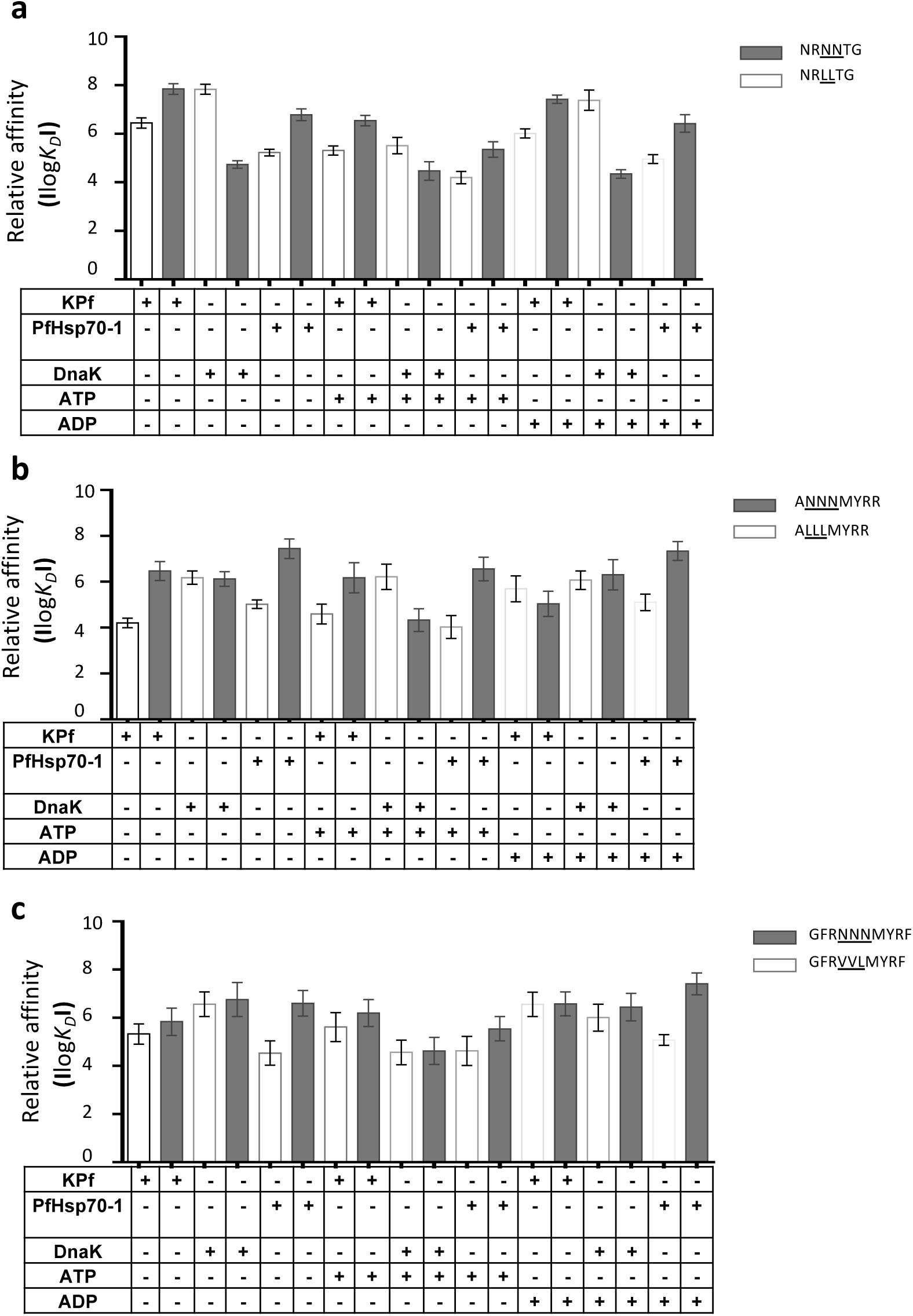
PfHsp70 and KPf preferentially bind to asparagine-enriched peptides. The relative affinities of DnaK, KPf and PfHsp70-1 for the various peptides were determined in the presence or absence of 5 mM ATP/ADP. Relative affinities for peptides: (**a**) NRLLTG versus NRNNTG (**b)** ALLLMYRR versus ANNNMYRR; (**c)** GFRVVLMYRF versus GFRNNNMYRF are depicted as bar graphs. The error bars shown were generated from three assays conducted using independent Hsp70 protein preparations. Statistical significance was determined by one-way ANOVA (*p*< *0.05*).

### SEC analysis of recombinant PfAdoMetDC protein co-produced with supplementary molecular chaperones

The recombinant PfAdoMetDC expressed in *E. coli* BL21 Star^™^ (DE3) cells endowed with endogenous levels of DnaK or supplementary chaperone sets was purified as previously described (Fig. S2; 29). Based on SEC analysis, monomeric PfAdoMetDC was eluted at the expected retention time at approximately 66 kDa based on MW calibrations used. Interestingly, PfAdoMetDC co-expressed with DnaJ and either DnaK/KPf/PfHsp70-1 eluted at a retention time around approximately 18 mins (Fig. 3a). On the other hand, recombinant PfAdoMetDC that was co-expressed with DnaK/KPf/PfHsp70-1+DnaJ+GroEL migrated at a slightly reduced pace suggesting that it assumed a more compact conformation than the protein expressed in the presence of only DnaJ-DnaK/KPf/PfHsp70-1 (Fig. 3a). Interestingly, the elution profile of PfAdoMetDC expressed in the absence of supplementary chaperones eluted at nearly the same retention time as the protein co-expressed with supplementary Hsp70 plus GroEL. PfAdoMetDC expressed in the absence of supplementary chaperones is known to aggregate and is hardly biochemically active (29, 49), and hence its comparatively long retention time could be due to its misfolded status Altogether, these findings suggest that supplementary GroEL may have improved folding of PfAdoMetDC towards a more compact conformation that was attained in the presence of DnaJ-DnaK/KPf/PfHsp70-1 chaperone sets.

**Figure 3.**
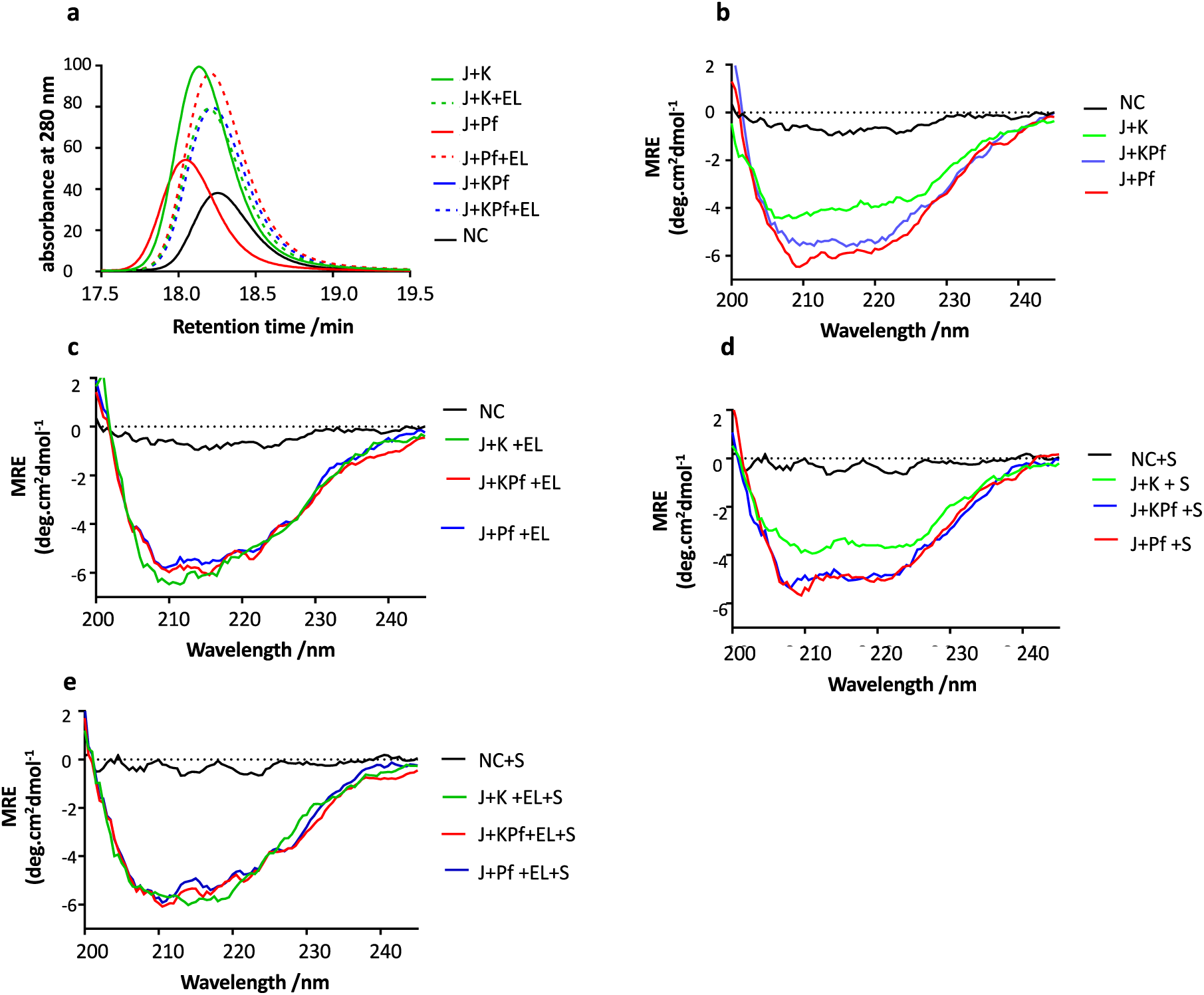
PfAdoMetDC co-produced with PfHsp70-1 and KPf is structurally unique secondary structure from that co-produced with DnaK. PfAdoMetDC was co-produced either alone (NC) or in the presence of supplementary chaperone sets: DnaK+DnaJ (J+K); DnaK+DnaJ+GroEL (J+K+EL); PfHsp70-1+DnaJ (J+Pf); PfHsp70+DnaJ+GroEL (J+Pf+EL); KPf+DnaJ (J+KPf); and KPf+DnaJ+GroEL (J+KPf+EL), respectively. The protein was then subjected to SEC-FPLC analysis (**a**); as well as far-UV CD spectrometric assays (**b-e**). CD spectrometric analysis was conducted either in the absence of substrate or in the presence of SAM (‘S’), the PfAdoMetDC substrate. The CD spectrum of PfAdoMetDC was presented as molar residue ellipticity [MRE] (deg.cm^2^dmol-1).

### PfAdoMetDC co-produced with PfHsp70-1 and KPf exhibits unique secondary structural features

We conducted CD spectroscopy in order to investigate the secondary structure of PfAdoMetDC co-expressed with the various chaperone sets. The assay was conducted at 22 °C and changes in mean residue ellipticity were monitored at wavelengths 200-240 nm (Fig. 3b, c). The assay was repeated in the presence of the SAM, the PfAdoMetDC substrate (Fig. 3d, e). Except for the control protein (produced without supplementary chaperones), PfAdoMetDC co-produced with supplementary DnaJ and any of DnaK/KPf/PfHsp70-1 exhibited two distinct negative troughs at 210 and 222 nm, respectively (Fig. 3b). The observed spectra are consistent with the predominantly ß-sheet fold of PfAdoMetDC (49). However, the trough for the spectra generated by PfAdoMetDC co-produced with KPf and PfHsp70-1 was deeper and distinct compared to that of PfAdoMetDC co-produced with DnaK. This suggests that PfAdoMetDC co-produced with supplementary DnaK lost some of its ß -sheet fold. Furthermore, PfAdoMetDC co-expressed with DnaJ+DnaK/KPf/PfHsp70-1 and supplementary GroEL generated a spectrum representing a strong ß-sheet fold (Fig. 3c). Notably, the inclusion of GroEL to the DnaJ-DnaK chaperone set, resulted in the production of PfAdoMetDC whose spectrum had a deeper trough than registered by PfAdoMetDC produced in the absence of supplementary GroEL. This suggests that the introduction of GroEL rescued the folding dilemma of PfAdoMetDC co-produced with DnaJ+DnaK (Fig. 3c). However, the folding of PfAdoMetDC co-produced with DnaJ+PfHsp70-1/KPf was not modulated by the introduction of supplementary GroEL, suggesting that KPf and PfHsp70-1 were sufficient for folding PfAdoMetDC.

Furthermore, the stability of the secondary structural fold of PfAdoMetDC was assessed in the presence of the substrate, SAM (Fig. 3d, e). The data obtained in the presence of SAM, mirrored that obtained in its absence, highlighting the consistency of the findings. Altogether, these findings suggest that while DnaK confounded PfAdoMetDC folding, the introduction of supplementary GroEL masked the protein folding deficiency of DnaK.

### Confirmation of PfAdoMetDC fold using ANS fluorescence-based assay

The ANS binds to hydrophobic (nonpolar) surfaces of proteins, through its nonpolar anilino-naphthalene group (50). For this reason, ANS is used to estimate the levels of exposed hydrophobic clusters of protein. In turn, the exposure of a protein’ s hydrophobic residues increases when a protein is misfolded. In the current study, the folded structures of PfAdoMetDC co-expressed with various combinations of chaperones were analysed using the ANS fluorescence spectra at wavelengths 425 - 600 nm (Fig. 4a, b). The assay was repeated in the presence SAM (Fig. 4c, d). PfAdoMetDC co-expressed with DnaJ+KPf or DnaJ+PfHsp70-1 exhibited the highest fluorescence signal at the same peak. On the other hand, PfAdoMetDC co-produced with DnaJ+DnaK, had a lower peak than that for protein co-produced with DnaJ+KPf or DnaJ+PfHsp70-1 (Fig. 4a). PfAdoMetDC expressed in the absence of supplementary chaperones had the lowest fluorescence signal (Fig. 4a). The introduction of SAM did not change the fluorescence spectra for PfAdoMetDC co-produced with DnaJ+KPf/PfHsp70-1/DnaK significantly (Fig. 4c). As we observed before, the addition of GroEL to the DnaJ+DnaK led to the generation of PfAdoMetDC whose fold was similar to that of the protein co-produced with DnaJ+KPf/PfHsp70-1+GroEL (Fig. 4b, d). This further suggests that PfAdoMetDC co-produced with DnaJ+DnaK required supplementary GroEL for complete folding. However, although the introduction of GroEL led to an increase in the fluorescence signal for PfAdoMetDC, it was also associated with a broad spectral peak which tended towards a blue shift (Fig. 4b, d). This suggests that PfAdoMetDC co-produced with supplementary DnaJ+DnaK+GroEL was still less folded than that co-produced with supplementary DnaJ+KPf/PfHsp70-1 with/without supplementary GroEL. This suggests that DnaK did not support PfAdoMetDC folding and while GroEL rescued misfolded PfAdoMetDC, it could not overcome all the folding deficiencies of PfAdoMetDC induced by co-expressing it with DnaK. Altogether, our findings demonstrated that PfAdoMetDC folding was effected in the following order, depending on the prevailing chaperone conditions:DnaJ+KPf/PfHsp70-1>DnaJ+DnaK>no supplementary chaperones. However, as previously noted, the introduction of supplementary GroEL to the DnaJ+DnaK combination,led to the production of PfAdoMetDC with improved fold.

**Figure 4.**
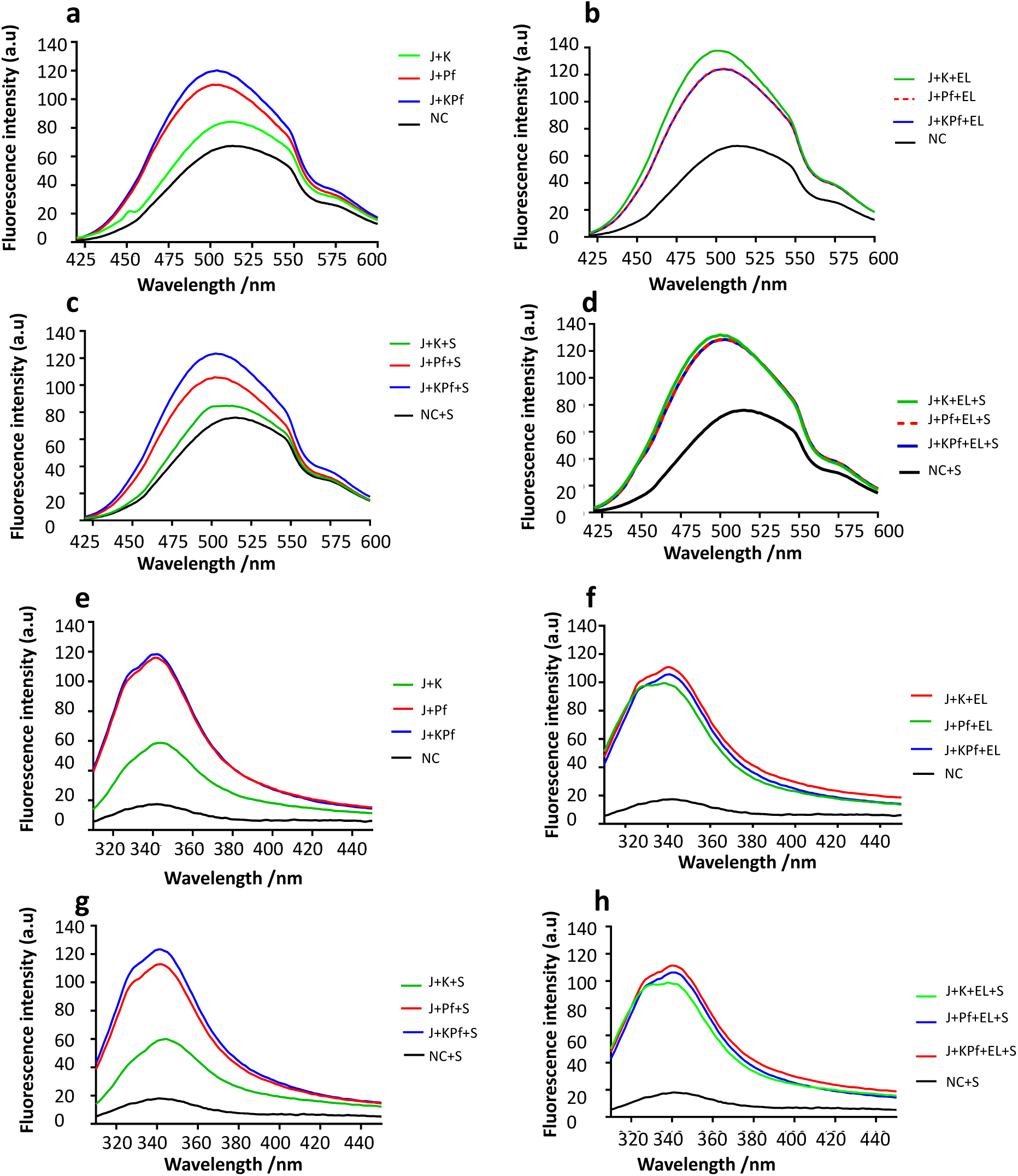
Co-production of PfAdoMetDC with PfHsp70-1 and KPf facilitates folding process. PfAdoMetDC co-expressed with the various chaperone sets was subjected to either ANS extrinsic fluorescence analysis (**a-d**) or intrinsic fluorescence analysis (**e-g**). PfAdoMetDC produced in the absence of supplementary chaperones (NC) or presence of various heterologous chaperones combinations: DnaK+DnaJ (J+K); DnaK+DnaJ+GroEL (J+K+EL); PfHsp70+DnaJ (J+Pf); PfHsp70+DnaJ+GroEL (J+Pf+EL); KPf+DnaJ (J+KPf); KPf+DnaJ+GroEL (J+KPf+EL). The fluorescence analysis was conducted in the absence or presence of the SAM (S), the PfAdometDC substrate.

### Analysis of tertiary structure of PfAdoMetDC using intrinsic tyrosine and tryptophan fluorescence

As a follow up study, the intrinsic fluorescence of PfAdoMetDC was analysed, taking advantage of the presence of a tryptophan residue (38W) located within the α-subunit of the protein. Due to its aromatic character, tryptophan is most often partially or fully buried in the hydrophobic core of protein. Upon disruption of the protein’ s tertiary structure, tryptophan becomes solvent exposed, leading to a blue shift in fluorescence (reviewed by 51). Tryptophan based fluorescence analysis of the tertiary structure was conducted by monitoring the emission spectra at 310 nm - 450 nm after an initial excitation at 295 nm as previously described (52, 53). As previously observed in the absence of supplementary GroEL, the intrinsic fluorescence signal intensity of PfAdoMetDC co-produced under the various chaperone conditions occurred in the following order: DnaJ+KPf>DnaJ+PfHsp70-1>DnaJ+DnaK>no supplementary chaperones (Fig. 4e). This trend remained the same upon repeating the assay in the presence of the substrate, SAM (Fig. 4f). PfAdoMetDC produced with supplementary DnaJ+DnaK in the absence of supplementary GroEL registered much less fluorescence intensity than the protein co-produced with either DnaJ+KPf or DnaJ+PfHsp70-1 (Fig. 4e-f). This further confirmed that PfAdoMetDC co-produced with supplementary DnaK assumed a unique conformation compared to PfAdoMetDC preparation that was co-produced with either KPf or PfHsp70-1. PfAdoMetDC that was co-produced with supplementary GroEL plus DnaK registered enhanced fluorescence signal, confirming that GroEL assisted PfAdoMetDC to overcome the folding snag it encountered in the presence of DnaK.

The assay was repeated to monitor the combined fluorescence signals of tryptophan and tyrosine as the protein possesses 36 tyrosine residues. The same trend was observed in which PfAdoMetDC produced in the absence of supplementary chaperones exhibited the lowest fluorescence signal, followed by PfAdoMetDC co-produced with DnaK chaperone system (Fig. S3). On the other hand, PfAdoMetDC co-expressed with either PfHsp70-1 or KPf exhibited the highest fluorescence signals, further suggesting that it had a unique conformation. However, the introduction of GroEL overcame the functional deficiencies of the DnaK system on PfAdoMetDC folding (Fig. S3). Analysis of PfAdoMetDC conformation was repeated in the presence of its substrate, SAM and the same trend was observed (Fig. S3), further confirming the consistency of the observed trend.

## DISCUSSION

PfHsp70-1 plays an important role in the survival and development of *P. falciparum*, the main agent of malaria. In addition, PfHsp70-1 is implicated in antimalarial drug resistance (reviewed in 5). It has been proposed that PfHsp70-1 constitutes a potential antimalarial drug target (54). Hsp70 chaperones are generally conserved, confounding their selectivity in drug targeting. However, there is growing evidence that notwithstanding their sequence conservation, these molecules exhibit specialised functional features across species. To the best of our knowledge this study for the first time demonstrates that PfHsp70-1 and its derivative, KPf, preferentially bound to asparagine enriched peptide substrates in vitro. Furthermore, expression of PfHsp70-1 and KPf in *E. coli* improved PfAdoMetDC folding. On the other hand, the enrichment of model Hsp70 peptide substrates with asparagine did not improve their affinity for *E. coli* DnaK. In addition, recombinant PfAdoMetDC folding did not benefit from co-production with supplementary DnaK in *E. coli*. Our findings do not only demonstrate the unique functional features between PfHsp70-1 and DnaK but also highlight the role of the SBD of Hsp70 in the interaction of this protein with nucleotide and the Hsp40 co-chaperone. One of the key features that sets apart PfHsp70-1 and other cytosolic Hsp70s of parasitic origin is the prominent presence of GGMP repeat motifs located in the C-terminal SBD of this protein (1, 6). The GGMP motif and other unique residues present in PfHsp70-1 may confer it with unique functional features.

In order to explore the functional specificity of PfHsp70-1, we systematically compared the structure-function features of PfHsp70-1 relative to those of DnaK, and their chimeric product, KPf. All the three proteins have previously been demonstrated to reverse the thermosensitivity of *E. coli* dnaK756 cells whose native DnaK is functionally compromised (15). The chimera, KPf is made up of the NBD of DnaK and shares the same SBD as PfHsp70-1. For this reason, KPf constituted an appropriate tool to explore the unique functional features of PfHsp70-1 relative to *E. coli* DnaK.

Our findings established that KPf exhibits structure-function features that are unique from its parental isotypes. In addition, co-expressing KPf, DnaK, and PfHsp70-1 with recombinant PfAdoMetDC in *E. coli*, led to the production of PfAdoMetDC protein with unique secondary and tertiary conformations. Our findings demonstrate that although the SBD regulates the functional specificity of PfHsp70-1, the cooperation of both the SBD and the NBD is important for this process. We established that KPf was nearly as stable to heat stress as PfHsp70-1, and that DnaK in turn was less stable than both KPf and PfHsp70-1 (Fig. 1). This suggests that the SBD of PfHsp70-1 which it shared with KPf, conferred stability to both KPf and PfHsp70-1. This is in concert with a previous study by Misra and Ramachandran (37) which reported that the SBD of PfHsp70-1 is important for the stability of the protein. As such, the SBD of DnaK may account for its comparatively lower stability to heat stress. Since the SBD of PfHsp70-1 and that of DnaK show some degree of sequence divergence (55), it remains to be established which structural motif accounts for the enhanced stability that the SBD of PfHsp70-1 confers to the protein. In agreement with these findings, we previously observed that the C-terminal EEVN residues of PfHsp70-x (*P. falciparum* Hsp70 that is exported to the parasite-infected red blood cell), contributes to the overall stability of the protein (30).

The structural features that KPf shared with PfHsp70-1 confirms the important role of the SBD of Hsp70 in modulating Hsp70 function. For example, both KPf and PfHsp70-1 were more stable to heat stress than DnaK. However, KPf also possesses key structure-function features that set it apart from PfHsp70-1. KPf exhibited much higher affinity for ATP than DnaK (>200-fold) and its affinity for ATP was at least an order of magnitude higher than of PfHsp70-1 (Table S2). Interestingly, the high affinity for ATP registered by KPf is mirrored by the fact that the chimeric protein was the most conformationally responsive to the presence of ATP (Fig. 1c). It is known that ATP and small molecule inhibitors that bind to the NBD modulate the global conformation of Hsp70 through allostery (15, 56, 57). Thus the enhanced conformational changes that ATP induced on KPf may be on account of the unique NBD:SBD interface of this chimeric protein. Interestingly, both KPf and PfHsp70-1 hydrolysed ATP more effectively than DnaK (Table S1). It has previously been reported that PfHsp70-1 exhibits higher ATPase activity than Hsp70s of human, bovine and *E. coli* origin (19). Here, we show that the SBD of Hsp70 plays an important role in regulating its ATPase activity.

PfHsp40 is a *P. falciparum* cytosol localised Hsp40 whose structure-function features resemble those of the canonical *E. coli* DnaJ/Hsp40 (20). PfHsp40 has been shown to stimulate the ATPase activities of cytosol-localised Hsp70s, including PfHsp70-1 and human Hsp70 (20). Here we demonstrated that PfHsp40 directly bound to all the three Hsp70s and exhibited comparative affinity for KPf and PfHsp70-1. However, its affinity for DnaK was an order of magnitude lower (Table S1). Furthermore, PfHsp40 stimulated the ATPase activity of KPf more effectively than it modulated the ATPase activities of either PfHsp70-1 or DnaK (Fig. 1d; Table S1). This finding suggests that the NBD of DnaK and the SBD of PfHsp70-1 constituting the domains of KPf, create a structurally unique NBD:SBD interface that promotes efficient hydrolysis of ATP in the presence of PfHsp40. Since the NBD of Hsp70 is highly conserved while its SBD is fairly divergent, the NBD:SBD interface of Hsp70 is regarded as a unique structural entity that regulates its functional specificity (6).

Some Hsp70s self-associate to form functional complexes that interact with co-chaperones to facilitate substrate refolding (58). A study reported that Hsp70 in its dimeric form interacts with Hsp40 to form a functional complex (45). We sought to interrogate the capability of all three proteins to self-associate. All the proteins were capable of self-association (Table 1). While KPf and PfHsp70-1 exhibited high affinity (nanomolar range in the presence of ATP), the self-association of DnaK was weaker (micromolar range) under similar conditions. These findings suggest that ATP promoted self-association of the three proteins and this is in line with a previous independent study which proposed that oligomerisation of DnaK is enhanced by ATP (45, 59). It is notable that both KPf and PfHsp70-1 exhibited comparably higher affinity for self-association than DnaK. It has been proposed that oligomerisation of Hsp70 is mediated by NBD:SBD interface and the linker segment (42, 58). Since both KPf and PfHsp70-1 share the same SBD and a highly conserved linker motif (60), these two subdomains that the two chaperones share may have accounted for their comparably higher propensity to form oligomers than DnaK.

It has been proposed that roughly 10% of *P. falciparum* proteome is marked by prion-like repeats and that more than 30% of the parasite proteins are characterized by glutamate/asparagine repeat segments (26). Furthermore, a previous study we conducted demonstrated that the red blood cell exported parasite Hsp70, PfHsp70-x, preferentially binds to peptides enriched with asparagine residues in vitro (30). For this reason, we explored the substrate binding preferences of PfHsp70-1 relative to KPf and DnaK. Three Hsp70 model peptides were employed in the study: the first, represented by the sequence, NRLLTG is a synthetic model substrate of DnaK (48); the second, ALLLMYRR is a derivative of chicken mitochondrial aspartate amino-transferase (7) while the third, GFRVVLMYRF, is a derivative of residues 256-268 of firefly luciferase (47).

As expected, in the presence of ADP, DnaK displayed high affinity (nanomolar range) for model Hsp70 substrate, NRLLTG. However, affinity of DnaK for the L-N substitution version of this peptide (NRNNTG), led to a drop in affinity (micromolar range). This is expected as DnaK prefers peptides enriched in hydrophobic residues (11). On the other hand, introduction of asparagine residues led both KPf and PfHsp70-1 to bind the peptide NRNNTG with higher affinity (nanomolar range) than they displayed for NRLLTG (Fig. 2; Table S3). Altogether, these findings show that whereas the SBD of DnaK preferred L residues to N residues in its substrate, L-N substitutions endeared the peptide for recognition by both PfHsp70-1 and KPf.

Similarly, the introduction of asparagine residues in the peptides, A<u>LLL</u>MYRR and GFRVVLMYRF, did not enhance DnaK’s affinity for the two peptides in the presence of ADP. However, both KPf and PfHsp70-1 bound to the peptide ANNNMYRR with higher affinity than they had for ALLLMYRR in the presence of ADP. PfHsp70-1 exhibited higher affinity for the peptide GFRNNNMYR than it exhibited for GFRVVLMYRF. However, while both DnaK and KPf exhibited high affinity (nanomolar range) for GRFVVLMYRF, the affinity of both chaperones for the asparagine enriched form of the peptide remained the same in the presence of ADP. Altogether these findings demonstrate that KPf and PfHsp70-1 bound to peptides of eukaryotic origin (ALLMYRR and GRFVVLMYRF) with comparable affinities. On the other hand, DnaK exhibited the highest affinity for its model substrate, NRLLTG. While, KPf and PfHsp70-1 exhibited general improved propensity to bind asparagine enriched forms of the peptides, the affinity of DnaK for the peptides was not enhanced by enriching the peptides with asparagine residues. This demonstrates that the SBDs of Hsp70s of plasmodial origin are generally tailored to bind asparagine rich peptides.

We previously demonstrated that PfAdoMetDC co-produced with supplementary DnaJ-DnaK was less active than protein co-produced with DnaJ-KPf/PfHsp70-1 (29). In the current study, we sought to establish whether co-expression of PfAdoMetDC with the various chaperone sets variably modulates its folded status. To this end, we purified recombinant PfAdoMetDC co-produced with the various chaperones we employed here and subjected it to SEC, CD and fluorescence spectrometric analyses. Our SEC analysis suggested that PfAdoMetDC that was co-expressed with DnaK/KPf/PfHsp70-1+DnaJ+GroEL was more compact than the protein expressed in the presence of only DnaJ-DnaK/KPf/PfHsp70-1 (Fig. 3). Thus supplementary GroEL appeared to marshal PfAdoMetDC folding towards a more compact conformation (Fig. 3). Interestingly, the elution profile of PfAdoMetDC expressed in the absence of supplementary chaperones eluted at nearly the same retention time as the protein co-expressed with the respective supplementary Hsp70 plus GroEL. The finding further suggests that the presence of supplementary GroEL modulated PfAdoMetDC to fold in a unique fashion.

The CD spectroscopic analysis confirmed that PfAdoMetDC is characterised by a dominant ß-sheet fold as previously reported (49). While the co-expression of PfAdoMetDC with DnaJ-KPf/PfHsp70-1 restored the conformation of the protein, co-production of the protein with DnaJ-DnaK led to partial loss of the ß -sheet fold of the protein (Fig. 3). This suggests that DnaK did not support PfAdoMetDC folding. However, combining GroEL with DnaJ-DnaK led to restoration of the ß-conformation of PfAdoMetDC (Fig. 3). While the introduction of GroEL appears to have modulated the fold of PfAdoMetDC co-produced with DnaK, it did not alter the apparent secondary structural fold of PfAdoMetDC co-produced with KPf/PfHsp70-1. This seems to suggest that the presence of KPf and PfHsp70-1 led PfAdoMetDC to a fully folded status. Subsequent analyses of PfAdoMetDC by ANS and the tryptophan/tyrosine fluorescence data corroborated that DnaK confounded PfAdoMetDC folding and that both KPf and PfHsp70-1 were more effective in facilitating its folding process (Fig. 4). Furthermore, based on CD-spectrometry, SEC analysis, and intrinsic fluorescence analyses (Fig. 4; Fig. S3), GroEL facilitated PfAdoMetDC to overcome the barriers presented by DnaK in its folding pathway. However, ANS-fluorescence analysis, revealed that although GroEL enhanced folding of PfAdoMetDC, the slight blue shift of the spectrum generated by the protein suggests that GroEL may not have fully folded PfAdoMetDC compared to the quality of protein co-produced with either KPf or PfHsp70-1.

By comparing the structure-function features of PfHsp70-1 relative to those of DnaK and the chimera, KPf, we established that the SBD of PfHsp70-1 is endeared to bind asparagine enriched peptides. This is quite an important attribute as *P. falciparum* proteome is fairly represented by glutamate/asparagine-rich candidates (26, 27). Given the propensity of such a proteome to misfold and aggregate, it seems the malaria parasite is endowed with an Hsp70 protein folding machinery that is primed to handle the protein folding demands of the parasite. This is particularly important during the development of malaria fever which would adversely impact on the parasite’s survival. That both PfHsp70-1 and its chimeric derivative, KPf, demonstrated capability to fold a plasmodial protein, PfAdoMetDC more effectively than *E. coli* DnaK further suggests that the architecture of Hsp70 of *P. falciparum* is biased to support folding of proteins of the malaria parasite. These findings are important for our understanding of the functional versatility of Hsp70 in spite of its apparent sequence conservation. In addition, the current findings contribute to ongoing efforts towards identifying small molecule inhibitors that selectively target Hsp70s of parasitic origin (56).

## EXPERIMENTAL PROCEDURES

### Expression and purification of recombinant molecular chaperones

Previously described plasmid constructs: pQE30/PfHsp70-1 (PlasmoDB accession number: PF3D7_0818900) encoding for PfHsp70-1 (61); pQE30/KPf encoding for KPf (29); pQE30/DnaK encoding for *E. coli* DnaK (15) were used for the expression of recombinant PfHsp70-1, KPf and DnaK proteins, respectively. In addition, a codon harmonized form of the *PfHsp40* gene (PlasmoDB accession number: PF3D7-1437900) was cloned into pQE30 (Qiagen, USA) using *Bam*HI and *Hind*III restriction. The DNA segment encoding PfHsp40 was produced by GenScript (USA) and integrity of the resultant pQE30/PfHsp40 was confirmed by agarose gel electrophoresis and DNA sequencing. The recombinant proteins were expressed in *E. coli* XL1 Blue cells in frame with N-terminally attached polyhistidine tag which facilitated purification using affinity chromatography as previously described (52). DnaK, PfHsp70-1 and KPf were successfully purified using sepharose nickel affinity chromatography under native conditions. PfHsp40 was purified as previously described (4). The production of the (His)_6_-tagged recombinant proteins was confirmed by Western analysis using mouse monoclonal α-His-horseradish peroxidase conjugated antibodies (Sigma Aldrich, USA). Imaging of the protein bands on the Western blot was conducted using the ECL kit (ThermoScientific, USA) as per manufacturer’s instructions. Images were captured using ChemiDoc Imaging System (Bio-Rad, USA).

### Co-expression of PfAdoMetDC with supplementary molecular chaperones

Next, we investigated the effect of KPf, PfHsp70-1 and DnaK on recombinant PfAdoMetDC folding upon expression in *E. coli*. This was conducted by co-expressing PfAdoMetDC in *E. coli* BL21 Star^™^ (DE3) cells with the following supplementary chaperone combinations: DnaJ-DnaK/PfHsp70-1/KPf plus/minus supplementary GroEL. To facilitate co-expression of PfAdoMetDC with DnaK + DnaJ, the pBB535 (encoding DnaK + DnaJ) and plasmid pASKIBA/PfAdoMetDC (a kind donation from Dr Bernd Bukau (Heidelberg University, Germany) were used to co-transform *E. coli* BL21 Star^™^ (DE3) cells. As previously described, the DnaK encoding segment was substituted by DNA segments encoding for either KPf or PfHsp70-1 cloned onto the pBB535 (29). Similarly, pBB542 (encoding for DnaK/PfHsp70/KPf + DnaJ + GroEL) were also used along with pASKIBA/PfAdoMetDC as previously described (29). Thus in summary, chemically competent *E. coli* BL21 Star^™^ (DE3) cells were co-transformed with pASKIBA/PfAdoMetDC along with either: pBB535 plasmid construct encoding for: PfHsp70-1 + DnaJ / KPf + DnaJ / DnaK + DnaJ or the plasmid pBB542 encoding for: PfHsp70-1 + DnaJ + GroEL / KPf + DnaJ + GroEL / DnaK + DnaJ + GroEL, respectively as previously described (29). Briefly, the transformed *E. coli* BL21 Star^™^ (DE3) cells were cultured in LB broth supplemented with 100 μg/ml ampicillin (Sigma Aldrich, Darmstadt Germany) to select for pASKIBA3 and 50 μg/ml spectinomycin (Sigma Aldrich, Darmstadt, Germany), to select for pBB535 and pBB542, respectively. As control, PfAdoMetDC expressed alone (without supplementary chaperones), was expressed in *E. coli* BL21 Star^™^ (DE3) cells as previously described (49). The expression of chaperones was initiated by addition of 1 mM IPTG and at OD_600_ = 0.2 while the expression of PfAdoMetDC was initiated by addition of 2 ng/ml anhydroxytetracycline (AHT) (IBA GmbH, Germany) at OD_600_ = 0.7. Cells were incubated for 24 hours (37 °C, 250 rpm) from the time of induction with IPTG. The cells were harvested by centrifugation (5000 g, 20 min, 4 °C) and the pellet was resuspended in lysis buffer (100 mM Tris pH 7.5; 300 mM NaCl; 10 mM Imidazole, containing 1 mM aminoethyl benzenesulfonyl fluoride hydrochloride (AEBSF) and 1 mg/ml of lysozyme). The recombinant PfAdoMetDC protein was purified using the Strep-Tactin Sepharose column (Invitrogen, California, USA) as described previously (29, 49). Purified PfAdoMetDC protein was quantified using the Bradford assay. Protein expression, solubility and purification were confirmed using SDS-PAGE analysis. Western blot analysis was used to verify the identity of PfAdoMetDC using monoclonal α-Strep-tag II antibodies (Novagen, Wisconsin, USA). Following Strep-Tactin Sepharose affinity column purification, PfAdoMetDC was analysed using an Äkta explorer system (Amersham Pharmacia Biotech, Pennsylvania, USA) fast protein liquid chromatography (FPLC), to determine migration profiles of the protein preparations that were co-expressed with the various chaperones sets. The migration pattern of PfAdoMetDC as determined by SEC was used to infer the compact/relaxed state of its fold. Retention times of the respective protein preparation were determined by SEC at 22 °C in running buffer (50 mM Tris-HCl, pH 7.5, 1 mM DTT, 500 mM NaCl and 0.02% [w/v] NaN_3_) using Bio-Select SEC 250-5 column (BioRad, California, USA) with a fractionation range of 1000 - 250 000 Da. The column was briefly equilibrated with the running buffer and calibrated using BioRad (BioRad California, USA) gel filtration standards (thyroglobulin [670 kDa], bovine gamma globulin [158 kDa], chicken ovalbumin [44 kDa], equine myoglobin [17 kDa], and vitamin B12 [1.35 kDa]), and the standard covered MW range of 1 350– 670 000 Da. The elution profiles of the protein preparations were determined as peaks obtained at 280 nm using a Spectra Series UV100 spectrophotometer (Thermo Fischer Scientific, Massachusetts, USA).

### Analysis of the secondary structure of the recombinant proteins

The secondary structure of the PfHsp70-1, DnaK and KPf protein was investigated using a far-UV circular dichroism (CD) (JASCO Ltd, UK) spectrometer as previously described (36, 62). Briefly, 0.2 μM of recombinant protein (DnaK, PfHsp70-1/KPf) in buffer PBS (137 mM NaCl, 2.7 mM KCL, 10 mM Na2HPO4, 2 mM KH2PO4, pH 7.5) was analysed and spectral readings monitored at 190 to 240 nm. Spectra was averaged for 15 scans after baseline correction (buffer without recombinant protein). The spectra were deconvoluted to α-helix, β-sheet, β-turn and unordered regions, using the Dichroweb server, (http://dichroweb.cryst.bbk.ac.uk; 63). The effect of heat stress on the secondary structure of the Hsp70s was investigated by monitoring the spectral changes of each protein at various temperatures (19 °C to 95 °C). The folded states of each Hsp70 protein at any given temperature was determined using spectral readings obtained at 222 nm as previously described (62). In a separate experiment, the analysis of PfAdoMetDC protein co-expressed with the various chaperone sets was conducted by monitoring the CD spectrum at 190 to 240 nm at 22 °C. The assay was repeated in the presence of PfAdoMetDC substrate, S-adenosylmethionine hydrochloride (SAM), as previously described with minor modifications (64). Briefly, PfAdoMetDC protein was preincubated in PBS supplemented with 100 μM SAM for 30 minutes prior to analysis at 25 °C. The resultant spectra were monitored and averaged following baseline correction (100 μM SAM using buffer void of the protein).

### Fluorescence based analysis of the tertiary structural organisation of the recombinant proteins

The tertiary structural conformations of DnaK, PfHsp70-1 and KPf proteins were assessed using tryptophan fluorescence as previously described (36). The recombinant proteins (0.45 μM) were incubated in buffer HKKM (25 mM HEPES-KOH pH 7.5, 100 mM KCl, 10 mM MgOAc) for 20 minutes at 20 °C. Fluorescence spectra were recorded between 300 nm and 400 nm after initial excitation at 295 nm using JASCO FP-6300 spectrofluorometer (JASCO, Tokyo, JAPAN). In addition, the effect of nucleotides on the conformation of each Hsp70 was investigated by repeating the assay in the absence or presence of 5 mM ATP/ADP. Similarly, biophysical characterization of PfAdoMetDC was conducted by monitoring both intrinsic (tryptophan and /tyrosine); and extrinsic (1-anilinonapthelene-8-sulfonate, ANS; Sigma Aldrich, Darmstadt, Germany) emission spectra as previously described (50). Since PfAdoMetDC possesses a single tryptophan residue and 33 tyrosine residues, the intrinsic PfAdoMetDC emission spectra was monitored at 300 - 400 nm. Furthermore, the extrinsic emission spectra generated upon ANS binding was conducted after an initial excitation at 370 nm of 2 μM PfAdoMetDC suspended in buffer HKKM supplemented with 100 μM ANS and monitoring the emission spectra at 400 nm – 600 nm. The assays were each repeated three times using independent batches of the respective protein.

### Evaluation of ATPase activities of DnaK, PfHsp70-1 and KPf

The basal ATPase activities of DnaK, PfHsp70-1 and KPf were evaluated based on the amount of released inorganic phosphate (*pi*) upon ATP hydrolysis as previously described (19, 62). Hsp70 proteins at final concentration of 0.4 μM were each incubated for 5 minutes in buffer HKMD (10 mM HEPES-KOH pH 7.5, 100 mM KCl, 2 mM MgCl2 and 0.5 mM DTT). The reaction was initiated by the addition of ATP at various concentrations (0 - 5 mM) and samples were analysed every 30 minutes for 4 hours. In order to determine the stimulatory effect of Hsp40 on the ATPase activity of Hsp70, the experiment was repeated in the presence of PfHsp40 (0.2 μM) as previously described (20).

### Determination of the nucleotide binding affinities of PfHsp70-1, DnaK and KPf

This assay was conducted at room temperature (25 °C) using a BioNavis Multi-p arametric surface plasmon resonance (MP-SPR; BioNavis, Tampere, Finland) spectroscopy system. PfHsp70-1/DnaK/KPf (as ligands) were each immobilised through amine coupling on the functionalized 3D carboxymethyl dextran sensors (CMD 3D 500L; BioNavis, Tampere, Finland). As analytes, aliquots of ATP/ADP were prepared at final concentration of 0, 1.25, 2.50, 5, 10 and 20 μM and were injected at 50 μl/min in each channel in series. Association was allowed for 2 minutes and dissociation was monitored for 8 min. Steady state equilibrium constant data was processed and analysed using TraceDrawer software version 1.8 (Ridgeview Instruments, Sweden).

### Investigation of self-association of Hsp70 proteins

Self-association of the recombinant proteins was determined using a MP-SPR (BioNavis, Tampere, Finland) as previously described (12, 61). As ligands, the recombinant Hsp70 proteins were immobilised on the CMD3D 500L chip. The analyte (respective Hsp70 protein) was prepared at various concentrations of 0, 125, 250, 500, 1000, 2000 nM, respectively and injected at 50 μl/sec onto each channel with immobilised ligands. A reference channel without immobilized protein served as control for non-specific binding and changes in refractive index. The analysis was conducted either in the absence or presence of 5 mM ATP/ADP. Association was allowed for 2 minutes, while dissociation was monitored for 8 minutes. Data analysis was conducted after double referencing by subtraction of both the baseline RU (buffer with ATP/ADP plus BSA as control protein) and RU from the reference channel. Kinetics steady-state equilibrium constant data was processed after concatenating the responses of all five analyte concentrations by global fitting using TraceDrawer software version 1.8 (Ridgeview Instruments, Sweden).

### Investigation of interaction of PfHsp40 with DnaK, PfHsp70-1 and KPf

The direct interaction of each Hsp70 with PfHsp40 was analysed using a MP-SPR (BioNavis, Tampere, Finland), as previously described. As ligands, recombinant DnaK, PfHsp70-1 and KPf were immobilised at concentrations of 1.0 μg/mL per each immobilization surface. The analyte (PfHsp40) was prepared at varied concentrations of 0, 125, 250, 500, 1000, 2000 nM and injected at 50 μl/sec onto the surface in series. The analysis was conducted both in the absence and presence of 5 mM of ATP/ADP. The ligand and analyte pairs were swapped, and data generated was analysed after global fitting. Association was allowed for 2 minutes, and dissociation was monitored for 8 minutes. Data analysis was conducted using TraceDrawer software version 1.8 (Ridgeview Instruments, Sweden).

### Interaction of DnaK, PfHsp70-1 and KPf with model peptide substrates

The respective affinity of each Hsp70 for model peptide substrate was investigated by MP-SPR. The Hsp70 ligands were immobilized as previously described (62). As analytes, the following model Hsp70 peptides were used: NRLLTG; NRNNTG; ALLLMYRR; ANNNMYRR; GFRVVLMYRF; and GFRNNNMYRF (30). The analytes were injected at varying concentrations (0, 125 nM, 250 nM, 500 nM, 1000 nM and 2000 nM) over the immobilized ligands (Hsp70s). The analytes were injected at a flow rate of 100 μl/ min and association and dissociation was allowed to occur for 10 minutes. The assay was conducted both in the absence and presence of 5 mM ADP/ATP. Analysis of association and dissociation data was conducted as previously described using TraceDrawer software version 1.8 (Ridgeview Instruments, Sweden).

## SUPPORTING DATA

**Table S1.** ATPase kinetics of DnaK, PfHsp70-1 and KPf.

| Proteins | $V_{max}$ (n/mol/min/mg) | $K_m$ (μM) |
| --- | --- | --- |
| DnaK | 14.33 (± 1.4) * | 179.1 (± 3.7) |
| DnaK+ PfHsp40 | 19.65 (± 0.81) | 354.6 (± 8.1) |
| PfHsp70-1 | 22.45 (± 1.8) | 370.8 (± 3.8) |
| PfHsp70-1+ PfHsp40 | 34.50 (± 0.62)* | 34.17 (± 0.5) |
| KPf | 27.6 (± 0.22) | 29.58 (± 2.2) |
| KPf + PfHsp40 | 42.6 (± 2.1)** | 25.65 (± 0.23) |

Table legends: The respective $V_{max}$ and $K_m$ values are shown. Standard deviations shown represent three independent assays made using separate protein purification batches. Statistical significance of data was validated by one way ANOVA test ( $p < 0.05$ \*) and ( $p < 0.01$ \*\*).

**Table S2.** Comparative ATP binding affinities for the Hsp70 proteins.

| Protein | ATP<br>$K_D/\mu$M (± standard deviation) |
| --- | --- |
| KPf | 0.38 (± 0.9) |
| PfHsp70-1 | 3.52 (± 1.1) |
| DnaK | 83.80 (± 6.6) |

The table shows equilibrium constant ( $K_D$ ) values representing the ATP binding affinities for DnaK, KPf, and PfHsp70-1. The ligand was the immobilized protein mounted onto HTE chip surface, while ATP represented the analyte which was injected at a flow rate of 100 μl/min. The standard deviations shown in parenthesis were obtained from three independent assays.

**Table S3.**
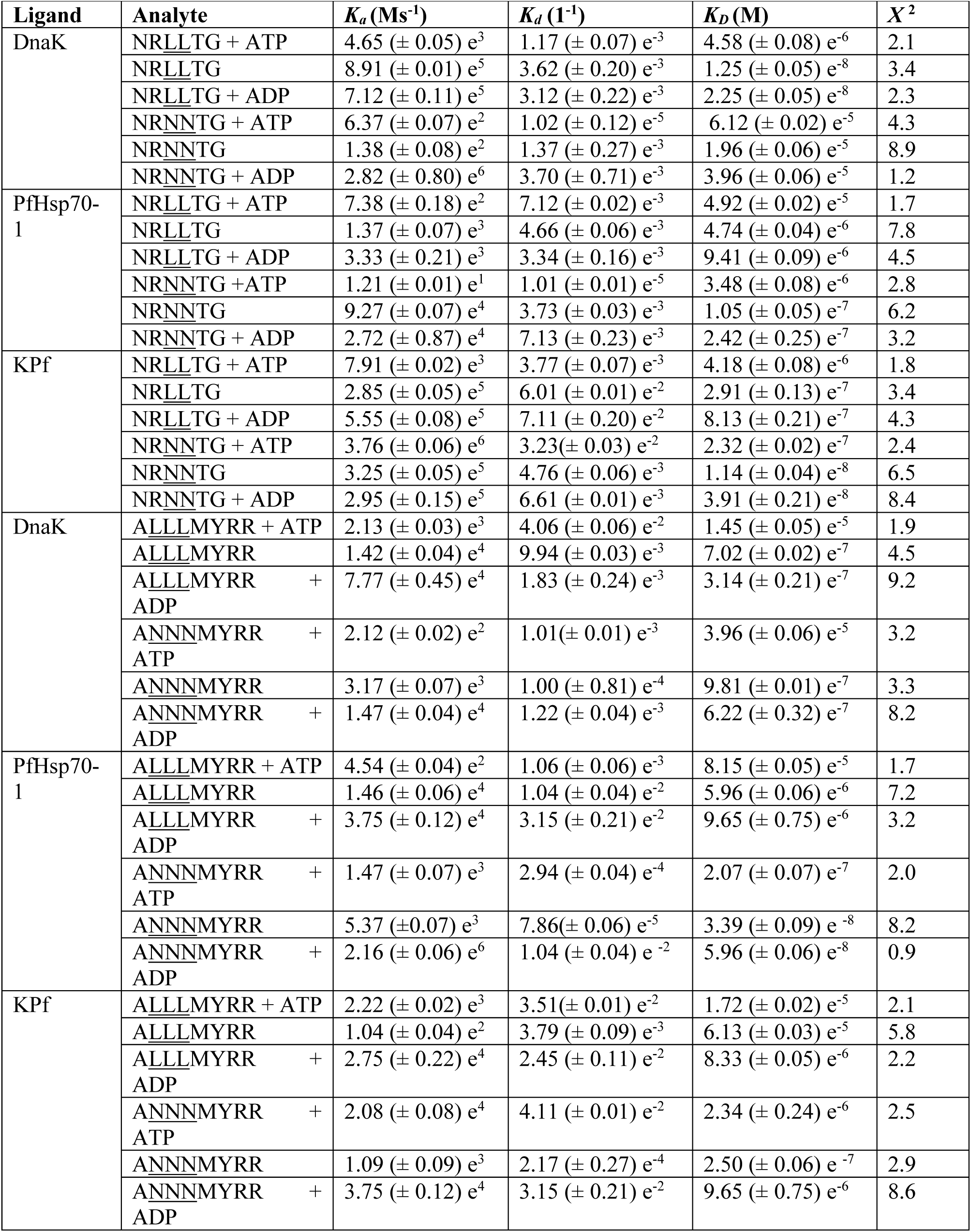
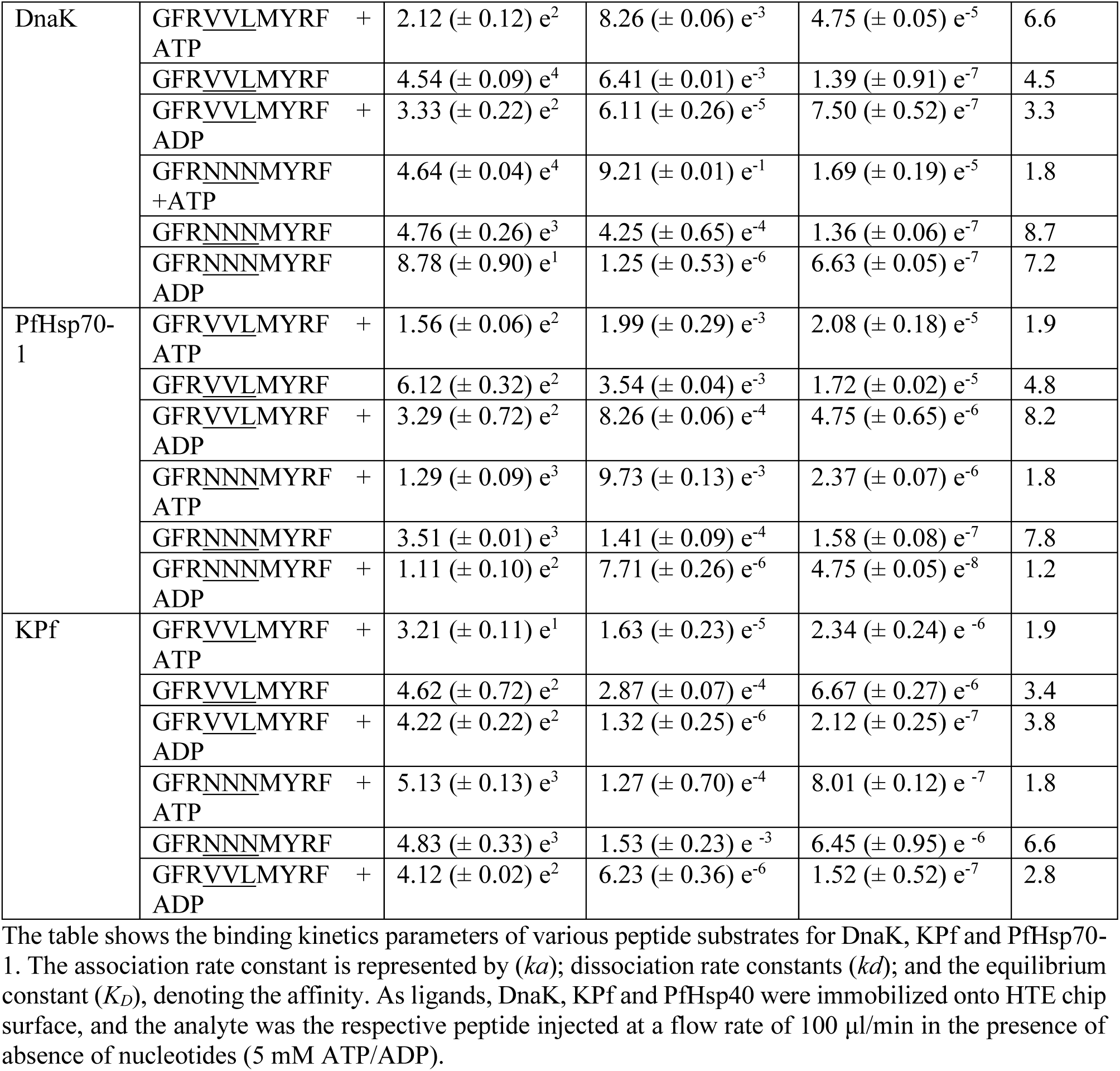
Kinetics interaction between DnaK, KPf and PfHsp70-1 with peptides.

| Ligand | Analyte | $K_a$ (Ms<sup>-1</sup>) | $K_d$ (1<sup>-1</sup>) | $K_D$ (M) | $\chi^2$ |
| --- | --- | --- | --- | --- | --- |
| DnaK | NRLLTG + ATP | 4.65 (± 0.05) e<sup>3</sup> | 1.17 (± 0.07) e<sup>-3</sup> | 4.58 (± 0.08) e<sup>-6</sup> | 2.1 |
|  | NRLLTG | 8.91 (± 0.01) e<sup>5</sup> | 3.62 (± 0.20) e<sup>-3</sup> | 1.25 (± 0.05) e<sup>-8</sup> | 3.4 |
|  | NRLLTG + ADP | 7.12 (± 0.11) e<sup>5</sup> | 3.12 (± 0.22) e<sup>-3</sup> | 2.25 (± 0.05) e<sup>-8</sup> | 2.3 |
|  | NRNNTG + ATP | 6.37 (± 0.07) e<sup>2</sup> | 1.02 (± 0.12) e<sup>-5</sup> | 6.12 (± 0.02) e<sup>-5</sup> | 4.3 |
|  | NRNNTG | 1.38 (± 0.08) e<sup>2</sup> | 1.37 (± 0.27) e<sup>-3</sup> | 1.96 (± 0.06) e<sup>-5</sup> | 8.9 |
|  | NRNNTG + ADP | 2.82 (± 0.80) e<sup>6</sup> | 3.70 (± 0.71) e<sup>-3</sup> | 3.96 (± 0.06) e<sup>-5</sup> | 1.2 |
| PfHsp70-1 | NRLLTG + ATP | 7.38 (± 0.18) e<sup>2</sup> | 7.12 (± 0.02) e<sup>-3</sup> | 4.92 (± 0.02) e<sup>-5</sup> | 1.7 |
|  | NRLLTG | 1.37 (± 0.07) e<sup>3</sup> | 4.66 (± 0.06) e<sup>-3</sup> | 4.74 (± 0.04) e<sup>-6</sup> | 7.8 |
|  | NRLLTG + ADP | 3.33 (± 0.21) e<sup>3</sup> | 3.34 (± 0.16) e<sup>-3</sup> | 9.41 (± 0.09) e<sup>-6</sup> | 4.5 |
|  | NRNNTG + ATP | 1.21 (± 0.01) e<sup>1</sup> | 1.01 (± 0.01) e<sup>-5</sup> | 3.48 (± 0.08) e<sup>-6</sup> | 2.8 |
|  | NRNNTG | 9.27 (± 0.07) e<sup>4</sup> | 3.73 (± 0.03) e<sup>-3</sup> | 1.05 (± 0.05) e<sup>-7</sup> | 6.2 |
|  | NRNNTG + ADP | 2.72 (± 0.87) e<sup>4</sup> | 7.13 (± 0.23) e<sup>-3</sup> | 2.42 (± 0.25) e<sup>-7</sup> | 3.2 |
| Kpf | NRLLTG + ATP | 7.91 (± 0.02) e<sup>3</sup> | 3.77 (± 0.07) e<sup>-3</sup> | 4.18 (± 0.08) e<sup>-6</sup> | 1.8 |
|  | NRLLTG | 2.85 (± 0.05) e<sup>5</sup> | 6.01 (± 0.01) e<sup>-2</sup> | 2.91 (± 0.13) e<sup>-7</sup> | 3.4 |
|  | NRLLTG + ADP | 5.55 (± 0.08) e<sup>5</sup> | 7.11 (± 0.20) e<sup>-2</sup> | 8.13 (± 0.21) e<sup>-7</sup> | 4.3 |
|  | NRNNTG + ATP | 3.76 (± 0.06) e<sup>6</sup> | 3.23 (± 0.03) e<sup>-2</sup> | 2.32 (± 0.02) e<sup>-7</sup> | 2.4 |
|  | NRNNTG | 3.25 (± 0.05) e<sup>5</sup> | 4.76 (± 0.06) e<sup>-3</sup> | 1.14 (± 0.04) e<sup>-8</sup> | 6.5 |
|  | NRNNTG + ADP | 2.95 (± 0.15) e<sup>5</sup> | 6.61 (± 0.01) e<sup>-3</sup> | 3.91 (± 0.21) e<sup>-8</sup> | 8.4 |
| DnaK | ALLLMYRR + ATP | 2.13 (± 0.03) e<sup>3</sup> | 4.06 (± 0.06) e<sup>-2</sup> | 1.45 (± 0.05) e<sup>-5</sup> | 1.9 |
|  | ALLLMYRR | 1.42 (± 0.04) e<sup>4</sup> | 9.94 (± 0.03) e<sup>-3</sup> | 7.02 (± 0.02) e<sup>-7</sup> | 4.5 |
|  | ALLLMYRR + ADP | 7.77 (± 0.45) e<sup>4</sup> | 1.83 (± 0.24) e<sup>-3</sup> | 3.14 (± 0.21) e<sup>-7</sup> | 9.2 |
|  | ANNNMYRR + ATP | 2.12 (± 0.02) e<sup>2</sup> | 1.01 (± 0.01) e<sup>-3</sup> | 3.96 (± 0.06) e<sup>-5</sup> | 3.2 |
|  | ANNNMYRR | 3.17 (± 0.07) e<sup>3</sup> | 1.00 (± 0.81) e<sup>-4</sup> | 9.81 (± 0.01) e<sup>-7</sup> | 3.3 |
|  | ANNNMYRR + ADP | 1.47 (± 0.04) e<sup>4</sup> | 1.22 (± 0.04) e<sup>-3</sup> | 6.22 (± 0.32) e<sup>-7</sup> | 8.2 |
| PfHsp70-1 | ALLLMYRR + ATP | 4.54 (± 0.04) e<sup>2</sup> | 1.06 (± 0.06) e<sup>-3</sup> | 8.15 (± 0.05) e<sup>-5</sup> | 1.7 |
|  | ALLLMYRR | 1.46 (± 0.06) e<sup>4</sup> | 1.04 (± 0.04) e<sup>-2</sup> | 5.96 (± 0.06) e<sup>-6</sup> | 7.2 |
|  | ALLLMYRR + ADP | 3.75 (± 0.12) e<sup>4</sup> | 3.15 (± 0.21) e<sup>-2</sup> | 9.65 (± 0.75) e<sup>-6</sup> | 3.2 |
|  | ANNNMYRR + ATP | 1.47 (± 0.07) e<sup>3</sup> | 2.94 (± 0.04) e<sup>-4</sup> | 2.07 (± 0.07) e<sup>-7</sup> | 2.0 |
|  | ANNNMYRR | 5.37 (± 0.07) e<sup>3</sup> | 7.86 (± 0.06) e<sup>-5</sup> | 3.39 (± 0.09) e<sup>-8</sup> | 8.2 |
|  | ANNNMYRR + ADP | 2.16 (± 0.06) e<sup>6</sup> | 1.04 (± 0.04) e<sup>-2</sup> | 5.96 (± 0.06) e<sup>-8</sup> | 0.9 |
| Kpf | ALLLMYRR + ATP | 2.22 (± 0.02) e<sup>3</sup> | 3.51 (± 0.01) e<sup>-2</sup> | 1.72 (± 0.02) e<sup>-5</sup> | 2.1 |
|  | ALLLMYRR | 1.04 (± 0.04) e<sup>2</sup> | 3.79 (± 0.09) e<sup>-3</sup> | 6.13 (± 0.03) e<sup>-5</sup> | 5.8 |
|  | ALLLMYRR + ADP | 2.75 (± 0.22) e<sup>4</sup> | 2.45 (± 0.11) e<sup>-2</sup> | 8.33 (± 0.05) e<sup>-6</sup> | 2.2 |
|  | ANNNMYRR + ATP | 2.08 (± 0.08) e<sup>4</sup> | 4.11 (± 0.01) e<sup>-2</sup> | 2.34 (± 0.24) e<sup>-6</sup> | 2.5 |
|  | ANNNMYRR | 1.09 (± 0.09) e<sup>3</sup> | 2.17 (± 0.27) e<sup>-4</sup> | 2.50 (± 0.06) e<sup>-7</sup> | 2.9 |
|  | ANNNMYRR + ADP | 3.75 (± 0.12) e<sup>4</sup> | 3.15 (± 0.21) e<sup>-2</sup> | 9.65 (± 0.75) e<sup>-6</sup> | 8.6 |
| DnaK | GFRVVLMYRF + ATP | $2.12 (\pm 0.12) e^2$ | $8.26 (\pm 0.06) e^{-3}$ | $4.75 (\pm 0.05) e^{-5}$ | 6.6 |
| | GFRVVLMYRF | $4.54 (\pm 0.09) e^4$ | $6.41 (\pm 0.01) e^{-3}$ | $1.39 (\pm 0.91) e^{-7}$ | 4.5 |
| | GFRVVLMYRF + ADP | $3.33 (\pm 0.22) e^2$ | $6.11 (\pm 0.26) e^{-5}$ | $7.50 (\pm 0.52) e^{-7}$ | 3.3 |
| | GFRNNNMYRF + ATP | $4.64 (\pm 0.04) e^4$ | $9.21 (\pm 0.01) e^{-1}$ | $1.69 (\pm 0.19) e^{-5}$ | 1.8 |
| | GFRNNNMYRF | $4.76 (\pm 0.26) e^3$ | $4.25 (\pm 0.65) e^{-4}$ | $1.36 (\pm 0.06) e^{-7}$ | 8.7 |
| | GFRNNNMYRF + ADP | $8.78 (\pm 0.90) e^1$ | $1.25 (\pm 0.53) e^{-6}$ | $6.63 (\pm 0.05) e^{-7}$ | 7.2 |
| PfHsp70-1 | GFRVVLMYRF + ATP | $1.56 (\pm 0.06) e^2$ | $1.99 (\pm 0.29) e^{-3}$ | $2.08 (\pm 0.18) e^{-5}$ | 1.9 |
| | GFRVVLMYRF | $6.12 (\pm 0.32) e^2$ | $3.54 (\pm 0.04) e^{-3}$ | $1.72 (\pm 0.02) e^{-5}$ | 4.8 |
| | GFRVVLMYRF + ADP | $3.29 (\pm 0.72) e^2$ | $8.26 (\pm 0.06) e^{-4}$ | $4.75 (\pm 0.65) e^{-6}$ | 8.2 |
| | GFRNNNMYRF + ATP | $1.29 (\pm 0.09) e^3$ | $9.73 (\pm 0.13) e^{-3}$ | $2.37 (\pm 0.07) e^{-6}$ | 1.8 |
| | GFRNNNMYRF | $3.51 (\pm 0.01) e^3$ | $1.41 (\pm 0.09) e^{-4}$ | $1.58 (\pm 0.08) e^{-7}$ | 7.8 |
| | GFRNNNMYRF + ADP | $1.11 (\pm 0.10) e^2$ | $7.71 (\pm 0.26) e^{-6}$ | $4.75 (\pm 0.05) e^{-8}$ | 1.2 |
| Kpf | GFRVVLMYRF + ATP | $3.21 (\pm 0.11) e^1$ | $1.63 (\pm 0.23) e^{-5}$ | $2.34 (\pm 0.24) e^{-6}$ | 1.9 |
| | GFRVVLMYRF | $4.62 (\pm 0.72) e^2$ | $2.87 (\pm 0.07) e^{-4}$ | $6.67 (\pm 0.27) e^{-6}$ | 3.4 |
| | GFRVVLMYRF + ADP | $4.22 (\pm 0.22) e^2$ | $1.32 (\pm 0.25) e^{-6}$ | $2.12 (\pm 0.25) e^{-7}$ | 3.8 |
| | GFRNNNMYRF + ATP | $5.13 (\pm 0.13) e^3$ | $1.27 (\pm 0.70) e^{-4}$ | $8.01 (\pm 0.12) e^{-7}$ | 1.8 |
| | GFRNNNMYRF | $4.83 (\pm 0.33) e^3$ | $1.53 (\pm 0.23) e^{-3}$ | $6.45 (\pm 0.95) e^{-6}$ | 6.6 |
| | GFRVVLMYRF + ADP | $4.12 (\pm 0.02) e^2$ | $6.23 (\pm 0.36) e^{-6}$ | $1.52 (\pm 0.52) e^{-7}$ | 2.8 |

The table shows the binding kinetics parameters of various peptide substrates for DnaK, Kpf and PfHsp70-1. The association rate constant is represented by ( $ka$ ); dissociation rate constants ( $kd$ ); and the equilibrium constant ( $K_D$ ), denoting the affinity. As ligands, DnaK, Kpf and PfHsp40 were immobilized onto HTE chip surface, and the analyte was the respective peptide injected at a flow rate of 100 μl/min in the presence of absence of nucleotides (5 mM ATP/ADP).

**Figure S1.**
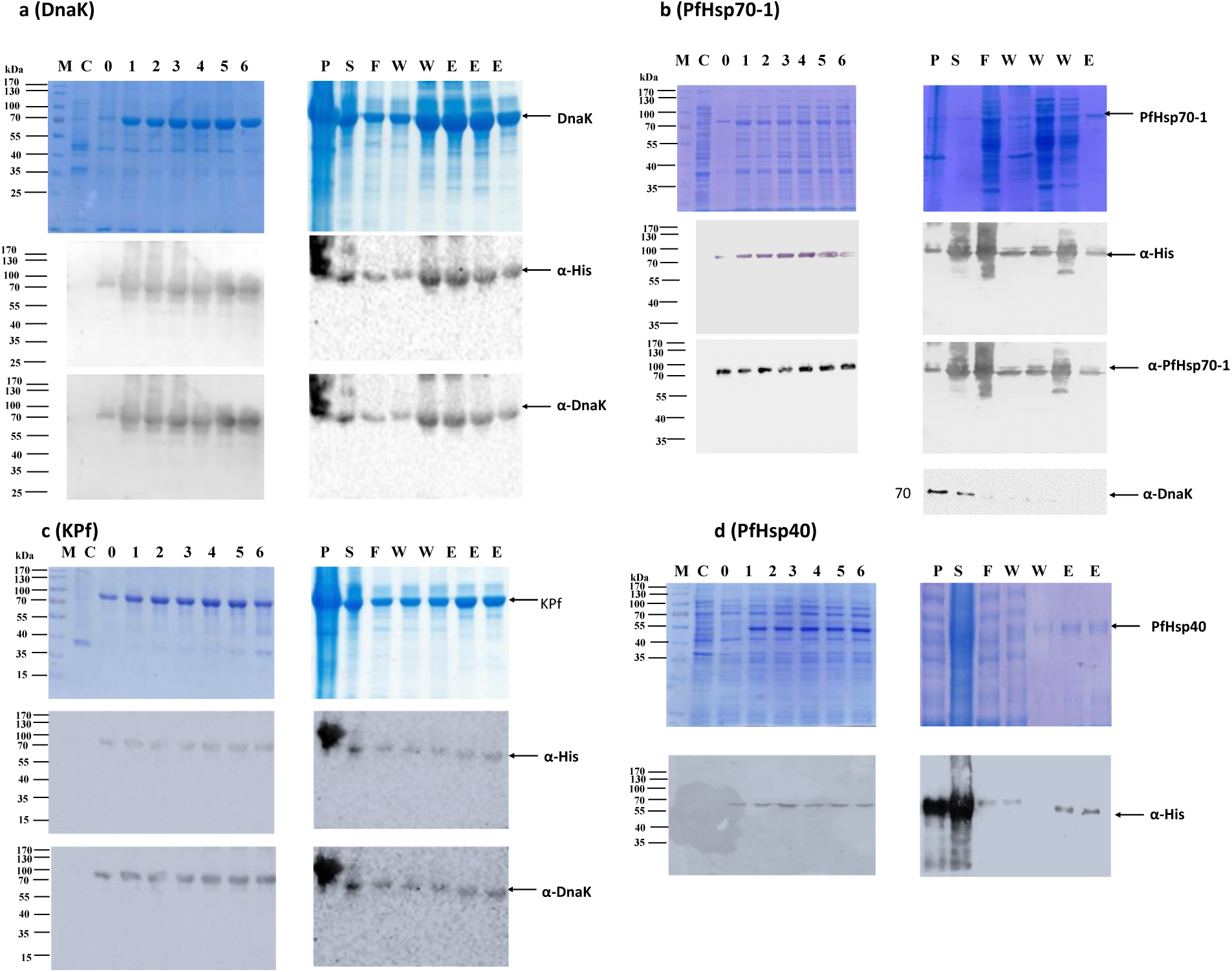
Expression and purification of recombinant proteins. The respective recombinant proteins were expressed in *E. coli* XL1 Blue cells, purified using affinity chromatography and were analysed with SDS-PAGE (12%): (**a**) DnaK; (**b**) PfHsp70-1; (**c**) KPf and (**d**) PfHsp40. Various lanes are denoted as follows: M– Page ruler (Thermo Scientific) in kDa; C—total extract for cells transformed with a neat pQE30 plasmid as control; 0– total extract of cells transformed with pQE30/PfHsp70-1/KPf/DnaK/PfHsp40 prior to IPTG induction; 1 - 6 - total cell lysate obtained hourly up to 6 h post induction; P, S; pellet and soluble fractions obtained from the total lysate of cells transformed with pQE30/PfHsp70-1/KPf/DnaK/PfHsp40, respectively; F-flow though, W-wash fraction; E-eluted fraction.

**Figure S2.**
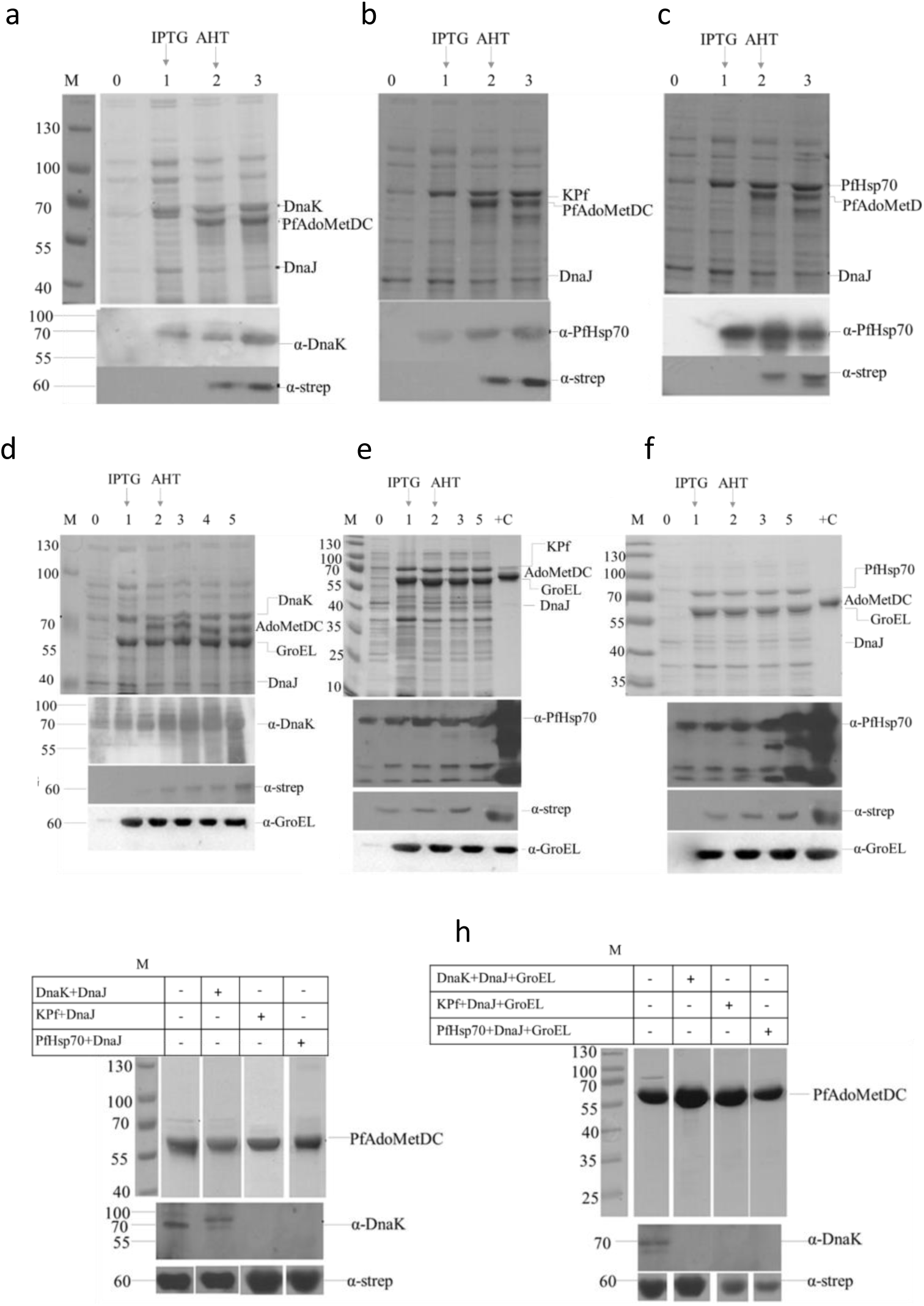
Co-expression and purification of PfAdoMetDC. The PfAdoMetDC recombinant proteins were co-expressed with different chaperone sets in *E. coli* BL21 Star (DE3) cells and were analysed by SDS-PAGE (12%) and Western blotting. Various co-expression chaperone sets were as follows: **(a)** DnaK+DnaJ; **(b)** KPf+DnaJ; **(c)** PfHsp70+DnaJ; **(d)** DnaK+DnaJ+GroEL; **(e)** KPf+DnaJ+GroEL; **(f)** PfHsp70+DnaJ+GroEL. Purification of PfAdoMetDC purified following expression with the various chaperone combinations is shown: (**g**); (**h**). Various lanes are denoted as follows: M– Page ruler (Thermo Scientific) in kDa; 0 – 5 total cell lysate obtained 0-5 hours IPTG induction; Note, induction with AHT was done 1 hour after IPTG induction. ‘’ +C’ ‘represents the positive control (purified PfAdoMetDC).

**Figure S3.**
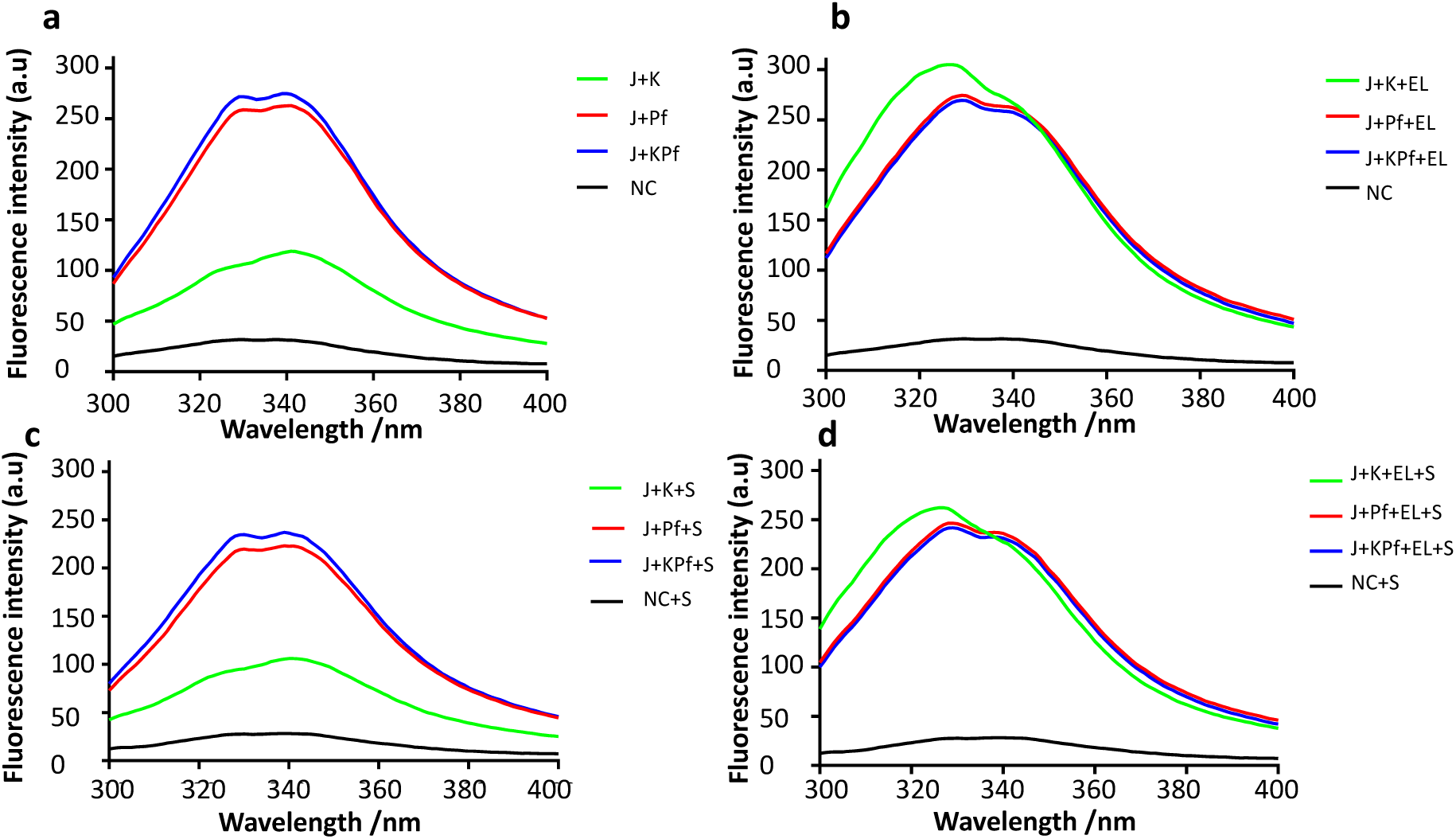
Combined tryptophan and tyrosine florescence spectra of PfAdoMetDC. Shown are the respective intrinsic combined tryptophan and tyrosine fluorescence signals of PfAdoMetDC co-expressed with: (**a**) Hsp70-DnaJ; (**b**) Hsp70-DnaJ-GroEL chaperone, respectively. PfAdoMetDC was analysed either in the absence of the substrate, SAM (**c**) and in the presence of SAM (**d**), respectively. The chaperone sets are denoted as: PfHsp70+DnaJ (J+Pf); PfHsp70+DnaJ+GroEL (J+Pf+EL); KPf+DnaJ (J+KPf); KPf+DnaJ+GroEL (J+KPf+EL). PfAdoMetDC produced in the absence of supplementary chaperones is represented by ‘NC’.

## FOOTNOTES

* This study was supported through a grant (L1/402/14-1) provided to AS by the Deutsche Forchungsgemeinshaft (DFG) under the theme, “German–African Cooperation Projects in Infectiology.” The authors are grateful to the Department of Science and Technology/National Research Foundation (NRF) of South Africa for providing an equipment grant (UID, 75464) and NRF mobility grant (UID, 92598) awarded to AS. CML is a recipient of the National Research Foundation Masters Scholarship; TZ is a recipient of the African-German Network of Excellence in Science junior researcher grant; AS is a recipient of a Georg Foster research fellowship awarded by the Alexander von Humboldt Foundation, Germany. We further wish to acknowledge Heini Dirr, retired Professor Emeritus, University of the Wistwatersrand, South Africa for technical support. The funders had no role in study design, data collection and analysis, and the decision to publish the findings.

1 Abbreviations used are: Hsp, Heat shock protein; NBD, nucleotide binding domain; SBD, substrate binding domain; KPf, *E. coli* DnaK-*Plasmodium falciparum* chimeric Hsp70; PfAdoMetDC, *Plasmodium falciparum* S-adenosylmethionine decarboxylase

